# Tumor-Initiating Cells Fine-tune the Plasticity of Neutrophils to Sculpt a Protective Niche

**DOI:** 10.1101/2025.04.04.647324

**Authors:** Weijie Guo, Jingyun Luan, Xuejie Huang, Daniel Leon, Jennifer Good, Benjamin Nicholson, Bijun Liu, Ama Owusu-Ofori, Evgeny Izumchenko, Ari J. Rosenberg, Nishant Agrawal, Breanna Bertacchi, Diana Bolotin, Matthias Gunzer, Iván Ballesteros, Andrés Hidalgo, Yuxuan ‘Phoenix’ Miao

## Abstract

The abundant accumulation of neutrophils in various solid cancers has been well recognized, but the functions of tumor-associated neutrophils (TANs) remain controversial. TANs have long been believed to be immune suppressive and have thus been referred to as “myeloid-derived suppressor cells”. However, effective tumor control induced by immunotherapy was recently found to be associated with strong neutrophil signatures. These seemingly contradictory findings highlight the unexpected degree of plasticity and heterogeneity unique to TANs. How the cellular plasticity and functional heterogeneity of TANs are regulated remains unknown. Here, we show that, while anti-PDL1/CD40 agonist immunotherapy can induce interferon responses to reprogram many TANs, allowing them to become plastic and regain anti-tumor activities in squamous cell carcinomas, a subset of TANs residing at the tumor-stroma interface can preserve their immune suppressive state. Importantly, by designing a reverse genetic screening, we identified a group of Sox2^Hi^ tumor-initiating cells (TICs) at the tumor-stroma interface that could upregulate Fatty Acid Desaturase 1 (Fads1) to produce arachidonic acid. This TIC-specific pathway can disrupt the interferon responsive potentials of TANs, preventing the interferon-mediated reprogramming. Thus, by fine-tuning the plasticity of neutrophils, TICs shape neutrophil heterogeneity and sculpt a protective micro-niche to survive from immunotherapy and drive cancer relapse.

## Main Text

Most solid tumors are complex ecosystems in which cancer cells closely interact with various cell types within the tumor microenvironment (TME) to drive tumorigenesis, metastasis, and immune evasion. Given the crucial roles of these communication networks, targeting these critical dialogues represents a promising therapeutic strategy to control malignancy. In particular, disrupting immune suppressive interactions through various immunotherapies holds great potential, as evidenced by the initial success of immune checkpoint blockade (ICB) therapy in many cancers (*1*). However, due to a lack of comprehensive understanding of these interactions between cancer cells and the TME, the effects of most immunotherapy treatments are not sustained, and many patients experience cancer relapse despite their initial responses (*2, 3*). Thus, an outstanding and puzzling question is how the dynamic spatial-temporal rearrangements of the TME induced by immunotherapy treatments are overcome by cancer cells to give rise to relapsed tumors.

The hierarchical organization of tumor populations add additional layers of complexity to the intertwined relationship between cancer cells and the TME (*4*). It is increasingly clear that many solid tumors, such as squamous cell carcinomas (SCCs) originating in various tissues, are initiated and maintained by a group of tumor-initiating cells (TICs) with strong stemness signatures (*5*). These TICs are known to activate special molecular programs to drive tumorigenesis and resistance to various therapies (*6, 7*). Importantly, recent studies have demonstrated that TICs can also survive robust anti-tumor immune responses evoked during immunotherapy which can otherwise clear differentiated cancer cells (*8, 9*). As a result, even though they still present targeted tumor antigens, these TICs can give rise to the relapsed tumors (*8–10*). These observations suggest that, compared to the rest of tumor populations, TICs must be equipped with unique immune resistance mechanisms. But how TICs acquire special protections during immunotherapy remain elusive.

Accumulating research investigating how normal tissue stem cells adapt to inflammation sheds key insights into understanding TIC-specific immune resistance. As long-lived cells, frequent exposure to inflammation could significantly impact the fitness of tissue stem cells (*11, 12*), debilitating their regenerative potential. Thus, stem cells residing in various barrier tissues are believed to be protected by an “immune privileged” niche, which is composed not only of physical barriers separating stem cells from infiltrating immune cells but also of a myriad of regulatory immune cells secreting various immune suppressive cytokines (*13, 14*). It is possible that TICs are shielded within a similar niche in the TME (*15*). However, our understanding of the composition, organization, and function of the special niche protecting TICs in various cancers is still unfolding.

Among various cell types in the TME that can interact with TICs, neutrophils are the least understood components. Comprehensive single-cell analysis of treatment free tumors has uncovered multiple cell states in TANs, especially a subset of dcTRAIL-R1^Hi^ CD101^+^CD14^Hi^ neutrophils, which were proposed to undergo deterministic and irreversible programming to acquire immune suppressive and pro-tumor functions (*16*). However, recent studies also revealed various anti-tumor functions of TANs. For example, TANs are shown to release elastase (*17*), nitric oxide (*18*), or reactive oxygen species (*19*) to directly kill cancer cells. Deep profiling has also unveiled that robust antigen presentation potential is frequently activated in a subset of TANs infiltrating various cancers (*20*),(*21*). Importantly, effective tumor control induced by immunotherapy was found to be associated with strong neutrophil signatures(*22, 23*). These seemingly contradictory findings suggested the unexpected degree of plasticity and heterogeneity unique to neutrophils infiltrating the tumors (*24*). Despite the progress, these recent findings have raised important new questions, such as whether the TANs are indeed plastic and can rapidly alter their cell states under different conditions, how effective immunotherapy impacts neutrophil plasticity and transitions between different cell states, how the spatial-temporal reprogramming of TANs contributes to tumor control, and how cancer cells overcome TAN-mediated anti-tumor immunity, leading to tumor relapse after immunotherapy treatments.

## Results

### Immunotherapies remodel the landscape and cell states of neutrophils in SCCs

To examine the plasticity of TANs, we employed a highly immunogenetic mouse spontaneous cancer model that was induced by exposing the mouse skin epithelium to chemical mutagen 7,12-dimethylbenz[*a*]anthracene (DMBA) followed by repeated applications of phorbol ester 12-O-tetradecanoylphorbol 13-acetate (TPA)(*25*) (Fig. 1A). This two-stage carcinogenesis process generates cutaneous SCCs bearing high mutational burdens which sensitize these tumors to immunotherapy treatments (*26*). Importantly, compared to previous studies which mostly profiled neutrophils infiltrating transplanted mouse tumors, the DMBA/TPA treatments induce autochthonous tumors whose developmental trajectory, heterogeneous composition, and complex ecosystem can better mimic human epithelial cancers. With this model, we next treated the mice bearing spontaneous skin SCCs with PDL1 blocking and CD40 agonist antibodies, which were recently demonstrated to induce anti-tumor activities in neutrophils (*23, 27*) (Fig. 1A). Immediately after the treatments (one day after the second dose), we collected these tumors and isolated total CD45+ immune cells. We subjected these cells as well as immune cells isolated from untreated tumors to single cell RNA-sequencing (scRNA-seq).

**Fig. 1.**
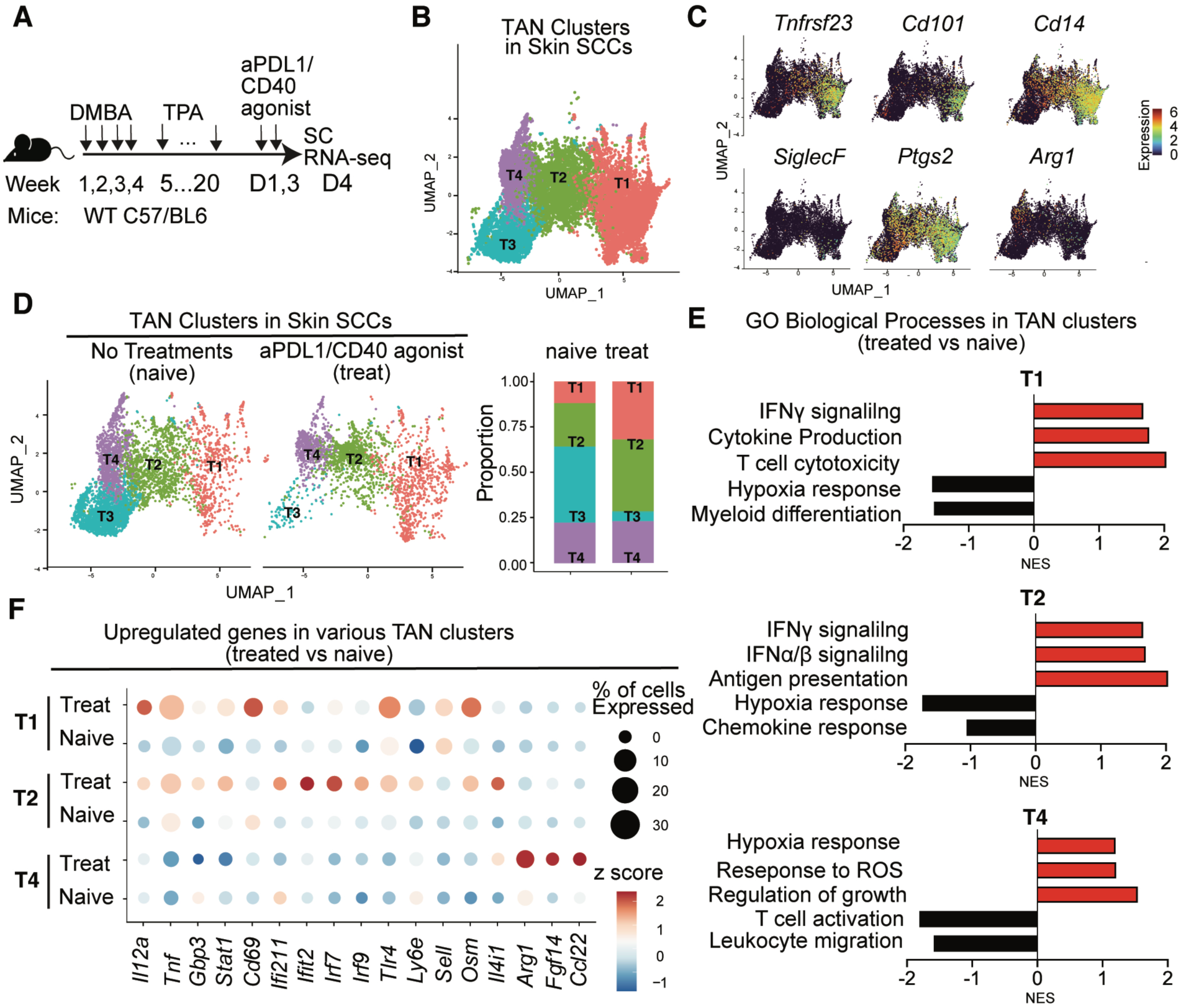
Immunotherapy induces distinct responses in different neutrophil subpopulations. **(A)** Schematic of experimental procedures for inducing spontaneous skin SCCs and analyzing the immune landscape changes induced by anti-PDL1/CD40 agonists immunotherapy. **(B and C)** UMAP showing the TAN subpopulation clustering in skin SCCs (B), and the signature or immune suppressive genes expressed (C) in various TAN clusters. **(D)** UMAP and stacked bar chart showing the changes in the composition of TAN subpopulations induced by the anti-PDL1/CD40 agonist treatments. **(E)** Up- or down-regulated pathways induced in each of the TAN clusters persisted after the anti-PDL1/CD40 agonist treatments. **(F)** Bubble heatmap showing transcripts of various immune stimulatory genes induced in cluster 1 and 2 (T1 and T2), and immune suppressive genes induced in cluster 4 (T4) upregulated after anti-PDL1/CD40 agonist treatments.

To profile the dynamic changes in neutrophils across various conditions, we adopted the split-pool combinatorial barcoding technique to perform scRNA-seq (*28*). This approach allows fixed cells to be collected from multiple conditions at different time points, and subsequently labeled, pooled, and sequenced simultaneously to minimize batch effects (*28*). With this method, we captured a comprehensive immune landscape in these tumors (fig. S1A). Using previously published neutrophil markers, such as *Mmp9, S100a8* or *Csf3r*, as reference to annotate cell identities, among these 40,500 immune cells sequenced, a total of 14,006 cells were identified as neutrophils (fig. S1B). Recent studies profiling pre-enriched neutrophils from grafted and treatment-free pancreatic cancer (*16*) or breast cancers (*29*) have identified several cell states of TANs. Our approach allowed us to identify similar subsets of neutrophils in skin SCCs without pre-enrichment (Fig. 1B). Consistent with previously published scRNA-seq results (*16, 29*), the cells in cluster 1 (T1) expressed *Tnfrsf23* (gene encoding dcTRAIL-R1) (*16*) and *SiglecF* (*23*), as well as high level of *Cd14* (*29*) and *Cd101* (*16*) (Fig. 1C), suggesting that this subset mainly contained mature neutrophils that were thought to undergo deterministic programming to acquire pro-tumor functions (*16, 29*). In contrast, *Tnfrsf23* could not be detected in cluster 4 (T4), and *Cd101* was also expressed at very low level in this cluster (Fig. 1C), suggesting these cells were immature neutrophils. Compared to grafted tumor models (*16*), spontaneous tumors appear to harbor more heterogeneous neutrophil populations. In particular, we identified additional transition states (T2 and T3) which still expressed major markers that could identify TANs with pro-tumor functions, such as *Tnfrsf23, Cd14*, or *Cd101,* but fewer cells did so when compared to T1 cluster (Fig. 1C). Each subset of TANs expressed specific signature genes (fig. S1C), and the cells in the transitional state shared certain markers with both T1 and T4, such as *Csf1* enriched in T1 and *Lpl* enriched in T4 (fig. S1C). Further supporting the developmental trajectory among these clusters, RNA velocity analysis confirmed that T1 cluster was the major mature subset, whereas the other three clusters developed towards these cells (fig. S1D). We then examined additional functional signatures of each subpopulation of TANs. Interestingly, in addition to sharing common immune suppressive mechanisms, such as expression of *Ptgs2* (*30*) (Fig. 1C), neutrophils in the T4 cluster expressed high level of *Arg1*, which could deplete arginine in the TME to block T cell functions (*31*) (Fig. 1C).

Next, we sought to investigate how immunotherapy treatments affect different TAN subpopulations. When we only focused on neutrophils and compared the representation of each TAN subpopulation before and after the treatments, we first found that neutrophils in the T3 cluster could be barely detected after injecting anti-PDL1 and CD40 agonist (Fig. 1D). This might be a result of their differentiation into more mature neutrophils in the T1 or T2 clusters, since both T1 and T2 populations expanded in treated tumors (Fig. 1D). In contrast, the T4 cluster persisted and maintained its representations (both cell number and percentage) after the treatments (Fig. 1D). We then performed pseudo-bulk analysis and profiled the gene expression changes within these persisting populations before and after immunotherapy treatment. To our surprise, although the dcTRAILR1^+^ and CD14^Hi^ neutrophils (both T1 and T2) were thought to be the major pro-tumor TANs (*16, 29*), after the immunotherapy treatment, the neutrophils in these cluster upregulated many pathways that could enhance anti-tumor immunity, such as genes involved in interferon responses, antigen presentation, and the regulation of T cell functions (Fig. 1E). However, compared to these TANs that appeared to undergo a conversion from being pro-tumor to anti-tumor TANs, these anti-tumor immune pathways were further downregulated in the T4 cluster (Fig. 1E). Instead, the TANs in the T4 cluster upregulated pathways involved in responding the reactive oxygen stresses and regulating tissue regeneration (Fig. 1E). Measuring the expression level of specific candidate genes confirmed these distinct responses between T1/2 and T4 TANs. It was reported that neutrophils with strong anti-tumor activity during immunotherapy treatments upregulate *Ly6e* (*22*) and *Sell* (*23*). Consistent with these results, the neutrophils in T1 and T2 appeared to be the major neutrophil population that upregulated *Ly6e* and *Sell*, as well as many other genes involved in anti-tumor immunity, such as *Il12a, Tnf*, and *Gbps* (Fig. 1F). In contrast, the neutrophils in T4 seemed to be resistant to these reprogramming effects induced during immunotherapy treatment (Fig. 1F). Instead, these TANs upregulated several metabolic immune checkpoint molecules, such as *Arg1* (*31*) and *Il4i1* (*32*), as well as factors that could recruit immune suppressive regulatory T cells, like *Ccl22* (*33*), or could directly promote tumor growth, such as *Fgf14* (*34*) (Fig. 1F).

### Immunotherapy-induced interferons reprogram a subset of TANs to gain anti-tumor functions

Given that the anti-PDL1/CD40 agonist treatments elicited distinct responses in different subsets of TANs, we sought to uncover the underlying mechanisms and to understand how different TAN subpopulations impacted the outcomes of immunotherapy treatments. Since more than 75% of TANs (both T1 and T2) upregulated genes involved in anti-tumor immunity and interferon responses (Fig. 1D-F), we first tested the significance of these changes. To achieve this goal, we generated *S100a8 Cre; R26-LSL-DTR* (Neu^DTR^) mice and grafted a syngeneic SCC line (PDVC57 cells) on either Cre+DTR+ mice or Cre-littermate controls. With these mice, we can deplete neutrophils in tumor-bearing mice at different time points when injecting diphtheria toxin (DT). Consistent with the known roles of TANs in suppressing anti-tumor immunity in mice when tumor-bearing mice did not receive any immunotherapy, when we depleted neutrophils in these mice (fig. S2A), the activities of tumor infiltrating T cells were enhanced (Fig. 2A). However, when DT was injected during the acute anti-PDL1/CD40 agonist treatments, neutrophil depletion significantly blunted anti-tumor activities in T cells (Fig. 2B). This result suggested that, while the majority of TANs were immune suppressive during the growth of primary tumors, many TANs became immune-stimulatory during anti-PDL1/CD40 agonist treatments. Further bolstering this conclusion, we found that after immunotherapy treatments, many TANs downregulated SiglecF, upregulated Ly6E, activated class II MHC (Fig. 2C), and all these molecular changes have been shown to closely associate with anti-tumor functions of TANs (*20, 22, 23*).

**Fig. 2.**
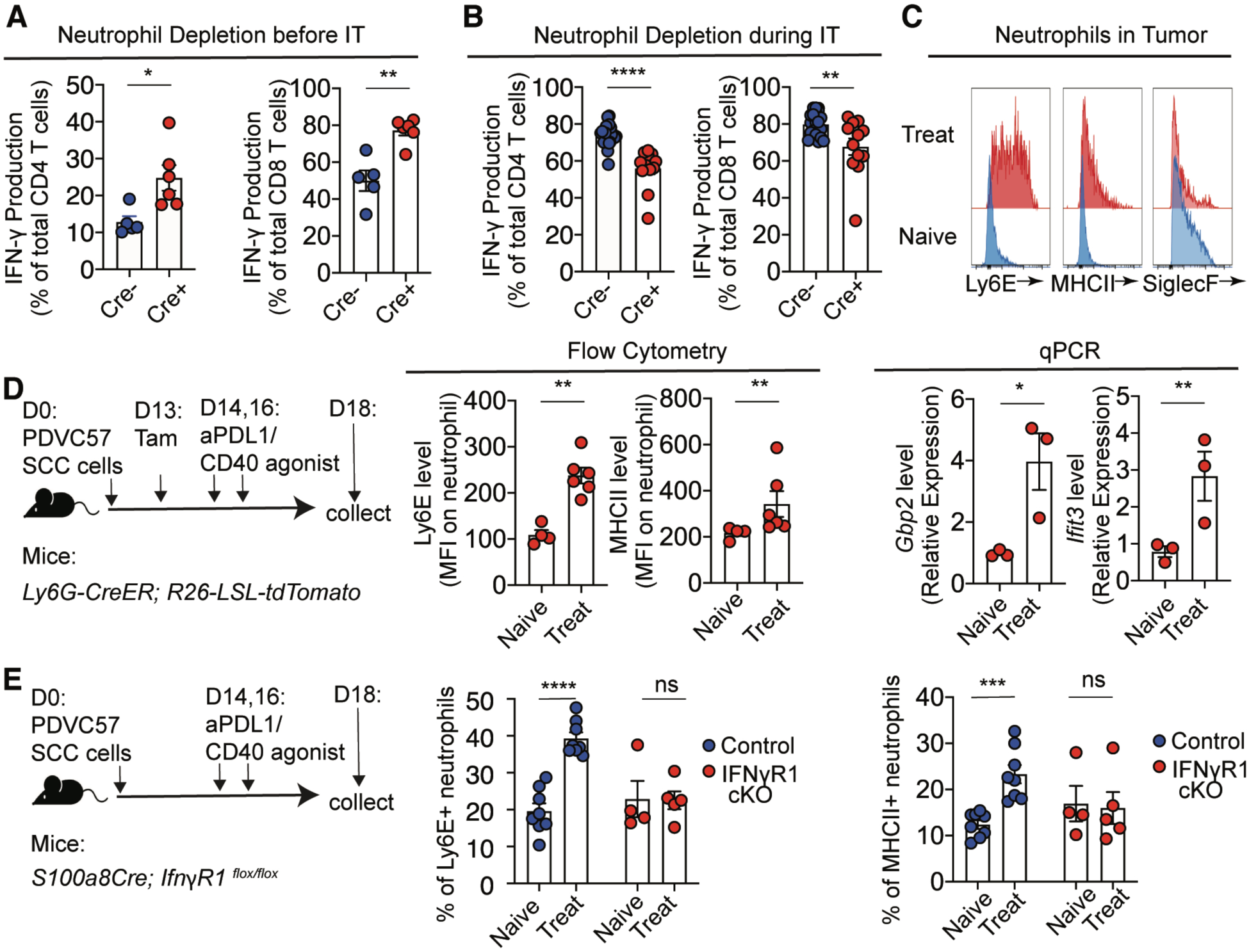
Immunotherapy-induced interferon response can reprogram TANs to gain anti-tumor functions. **(A)** Flow cytometry quantification of T cell responses when TANs were depleted before any immunotherapy (IT) treatments. n=5 for Cre- and n=6 for Cre+ group. **(B)** Flow cytometry quantification of T cell responses when TANs were depleted during the anti-PDL1/CD40 agonist treatments. n=15 for Cre- and n=12 for Cre+ group. **(C)** Representative flow cytometry histogram showing the upregulation of markers associated with anti-tumor activity (e.g. Ly6E, MHCII) or downregulation of markers associated with pro-tumor functions (e.g. SiglecF). **(D)** Experimental scheme, flow cytometry quantification of Ly6E and MHCII expression (left) and qPCR (right) quantification of interferon stimulated gene (*Gbp2* and *Ifit3*) expression in lineage traced pre-existing TANs (Tomato+) before or after the immunotherapy treatment. n=4 for naïve and n= 6 for treated group in flow cytometry, n=3 for each group in qPCR. **(E)** Experimental scheme and flow cytometry quantification of Ly6E or MHCII expression in neutrophils when the IFNγ receptor is deleted specifically in neutrophils (IFNγR1 cKO). n=8 for both naïve and treated control group, n=4 for naïve and n=5 for treated cKO group. Graphs show representative results from one of the three repeats for each experiment and was presented as mean ± SEM. *p < 0.05; **p < 0.01; ***p < 0.001; ns, non-significant.

The shift in neutrophil functions during the anti-PDL1/CD40 agonist treatments could be a result of biogenesis of new anti-tumor neutrophils in the bone marrow that were induced by immunotherapy treatments and recruited to the tumors, replacing the original suppressive TANs in the TME. However, analyzing the phenotypes of neutrophils in the bone marrow or spleen on the same tumor-bearing mice suggested that this might not be the case since the phenotype conversion in TANs were not observed in neutrophils outside the TME (fig. S2B). To further test whether immunotherapy treatment can convert immune-suppressive TANs into anti-tumor neutrophils, we designed a lineage tracing assay to distinguish the original TANs that were present in the TME before treatments from any newly generated neutrophils in the bone marrow after receiving anti-PDL1/CD40 agonists (Fig. 2D). Driven by this goal, we grafted PDVC57 SCC cells on the *Ly6G-CreER; R26-LSL-tdTomato* mice that could genetically label the neutrophils with Tomato upon receiving tamoxifen (*35*) (Fig. 2D). After primary tumor formation, we injected one dose of tamoxifen and confirmed that at least 10% of TANs could be labelled with Tomato (fig. S2C). Following genetic labeling of pre-existing TANs, we treated these lineage-tracing mice with anti-PDL1/CD40 agonist and then analyzed Tomato+ neutrophils. Both flow cytometry and qPCR analysis of sorted Tomato+ TANs confirmed that these Tomato+ neutrophils effectively acquired markers associated with anti-tumor immunity (e.g. Ly6E or MHCII) and activated interferon-regulated genes (*e.g. Gbp2* or *Ifit3*; Fig. 2D). This finding confirmed our speculation that the TANs that were already present within the TME became plastic and could be directly reprogrammed to gain anti-tumor functions. This finding is surprising because the anti-tumor functions appeared to be mainly activated within the subset of dcTRAILR1^Hi^ CD14^Hi^ neutrophils (Fig. 1C,D), which were considered the major population undergoing irreversible programming to mediate pro-tumor functions (*16, 29*). To further confirm that these TANs can be plastic and reprogrammed to re-gain anti-tumor activities, we employed fluorescence-activated cell sorting (FACS) to isolate the CD14^Hi^, CD14^Mid^, and CD14^Low^ TANs from untreated naive tumors. We then treated these different subsets of TANs with combined IFNα and IFNγ in culture (fig. S2D). Interestingly, we found that MHCII and Ly6E could be most efficiently induced in the CD14^Hi^ cells (fig. S2E,F). To further gain the molecular insight into how these pro-tumor TANs can be directly reprogrammed to regain anti-tumor activities during immunotherapy treatments, we generated *S100a8-Cre; IfnγR1 ^flox/flox^*(IFNγR1 cKO) mice to specifically block the interferon responses in neutrophils. Upon anti-PDL1/CD40 agonist treatments, we found that ablation of IFNγ signaling specifically in neutrophils abolished these neutrophil responses. As a result, TANs maintained their pro-tumor phenotypes and failed to upregulate Ly6E or class II MHC (Fig. 2E).

### Distinct spatial distributions of neutrophil responses during immunotherapy treatment

These lineage tracing and genetic manipulation experiments confirmed our conclusion from the single cell analysis that interferons can enhance the plasticity of many TANs, allowing these myeloid cells to regain immune stimulatory potentials during immunotherapy treatments. At the same time, these findings also triggered our interest in exploring why and how a small fraction of TANs (T4 cluster) could resist these reprogramming effects to preserve or even enhance their pro-tumor functions after the immunotherapy treatments (Fig. 1C,D). The distinct phenotypes and functional states displayed by different subsets of TANs prompted us to hypothesize that these neutrophil subpopulations might be positioned in separate regions in the TME, and their spatial distribution could dictate their differential responses to immunotherapies. To test this idea, we employed GeoMx spatial transcriptomics analysis to determine the distribution patterns of various neutrophil cell states in the spontaneous mouse SCCs. Briefly, we again utilized DMBA/TPA to induce autochthonous cutaneous SCCs on the neutrophil reporter ‘Catchup^IVM-red^’ (*Ly6G-Cre; R26-LSL-tdTomato*) mice (*36*) followed by anti-PDL1/CD40 agonist treatments. Next, we subjected the thin sections collected from both naïve and immunotherapy-treated tumors to the GeoMx spatial transcriptome profiling (Fig. 3A). This approach allowed us to combine immunofluorescence staining with digital barcoding to perform multiplexed and spatially resolved profiling of the gene signatures in neutrophils accumulated at different locations in the TME. When using Tomato signals and Keratin 14 antibody to visualize neutrophils and cancer cells respectively, we identified that most TANs were located either within the stroma (stromal TANs) or at the tumor-stroma interface (interface TANs, Fig. 3A). Thus, we selected Regions of Interests (ROI) within these two areas from tumors on naïve mice or animals receiving anti-PDL1/CD40 agonist. We then sequenced the Tomato+ neutrophils from selected ROIs and mapped their gene expression pattern to their original locations. Interestingly, we found that, similar to the T1 and T2 clusters, the neutrophils in the stroma exhibited global upregulation of interferon responses and other anti-tumor immunity-related pathways (Fig. 3B,C). In contrast, neutrophils located at the tumor-stroma interface showed downregulated interferon responses compared to those in the stroma (Fig. 3B,C).

**Fig. 3.**
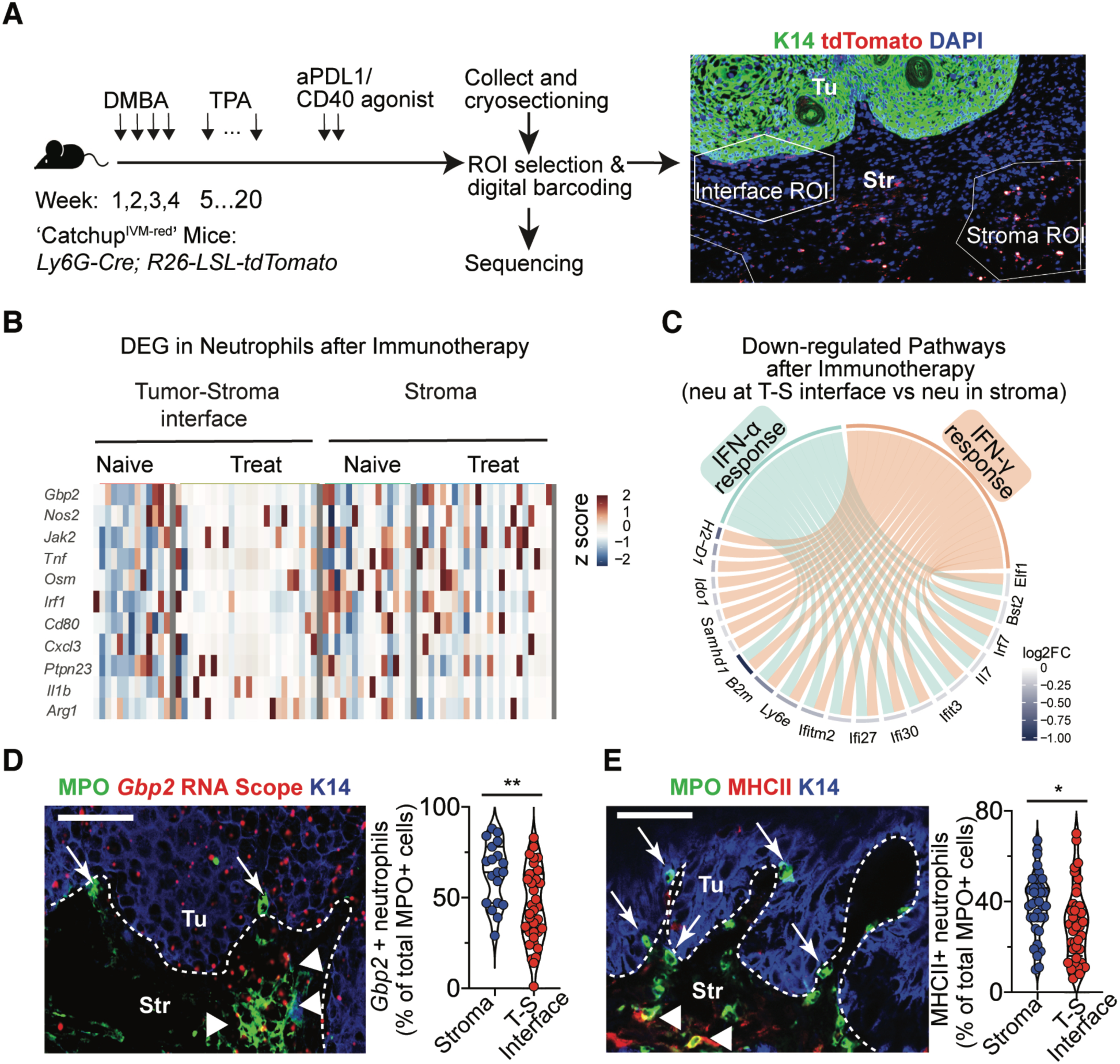
Spatial distribution of distinct neutrophils responses during immunotherapy treatments. **(A)** Experimental scheme and representative IF images of skin SCCs collected for selecting different ROIs and spatial transcriptomic analysis. **(B)** Heatmap showing scaled expression of anti-tumor immunity-related genes in neutrophils located at the tumor-stroma interface or in stroma before and after anti-PDL1/CD40 agonist treatments. **(C)** Chord diagram showing the down-regulated pathways in neutrophils located at the tumor-stroma interface compared to the neutrophils in stroma during immunotherapy treatments. **(D and E)** Representative IF images and quantifications of *Gbp2* (D) or MHCII (E) expression in neutrophils among total TANs located at the tumor-stroma interface (arrow) or in stroma (arrowhead) after immunotherapy treatments. The *Gbp2* mRNA was stained with RNAscope probes and neutrophils were stained with MPO antibody. K14 staining was used to depict tumor regions, and the tumor boarder was labelled with white dotted line. Scale bars: 50μm. *p < 0.05, **p < 0.01.

These results were intriguing and provided us preliminary insights. To further validate the pattern of interferon responses in TANs revealed by spatial transcriptome profiling, we conducted additional analyses. First, we used RNAscope-based imaging to visualize the spatial organization of interferon responses in TANs during anti-PDL1/CD40 agonist treatments. This experiment confirmed that interferon-stimulated genes (e.g. *Gbp2*) were strongly induced in TANs expanding in the stroma after the anti-PDL1/CD40 agonist treatments, whereas most neutrophils residing at the tumor-stroma interface were less responsive (Fig. 3D). We further confirmed this spatial pattern of anti-tumor responses in TANs through immunofluorescent staining for MHCII (Fig. 3E). Notably, other cell types at the tumor-stroma interface, including many cancer cells, were still strongly labelled with *Gbp2* probes (Fig. 3D), indicating that the deficient interferon response in neutrophils was not due to a lack of IFNγ production in the tumor-stroma interface. Collectively, these results suggest that the tumor-stroma interface represents a unique micro-niche that can actively block a subset of neutrophils from responding to immunotherapy-induced interferons, blunting their plasticity. Consequently, similar to the TANs from IFNγR cKO mice (Fig. 2E), the subset of neutrophils within this micro-niche remains to be immune suppressive during immunotherapy treatments.

### Sox2 amplification endows cancer cells with the capacity to limit the plasticity of TANs by blunting their interferon responses

Next, we sought to uncover the mechanisms specifically activated at the tumor-stroma interface that were responsible for blocking the TANs from gaining plasticity at this site. The tumor-stroma interface in SCCs is an interesting location, as it is where a group of TGFβ-responsive TICs accumulate (*37, 38*). Importantly, these cells were shown to be able to survive robust immunotherapy treatment and give rise to relapsed tumors without losing the targeted tumor antigens (*9*). Thus, we hypothesized that, in order to survive from immunotherapy treatments, TICs were equipped with a unique molecular program to modulate TANs. Given that most TIC functions are activated by stem cell-specific transcription factors (TFs), we tested our hypothesis by designing a reverse genetic screening with the goal of determining the key TIC-specific TFs that can endow cancer cells with the capacity to block TANs from responding to anti-PDL1/CD40 agonist treatments. Briefly, by isolating TICs from DMBA/TPA induced skin SCCs for bulk RNA-seq, we were able to compare the transcriptome of different tumor populations and identify a cohort of TFs that were specifically enriched in TICs (Fig. 4A). We then individually amplified these TFs in the immunogenic SCC cells (PDVC57) and grafted them on WT mice (Fig. 4A). Following anti-PDL1/CD40 agonist treatments, we analyzed the phenotypes of TANs infiltrating various SCC tumors overexpressing different TIC-specific TFs. This strategy allowed us to identify that, when compared to TANs isolated from the tumors formed by the SCC cells amplifying control vectors or most other tested TIC-specific TFs, only in tumors derived from SCC cells where Sox2 was amplified (Sox2^Hi^), TANs maintained the CD14^low^ Ly6E^low^ phenotypes (Fig. 4A). When TANs respond to anti-PDL1/CD40 agonist, they activate class II MHC (*22*). As we expected, this critical molecular change could not be induced in TANs when their surrounding cancer cells expressed high level of Sox2 (Fig. 4B). These results further confirmed the immune modulatory effects of Sox2^Hi^ cancer cells on TANs.

**Fig. 4.**
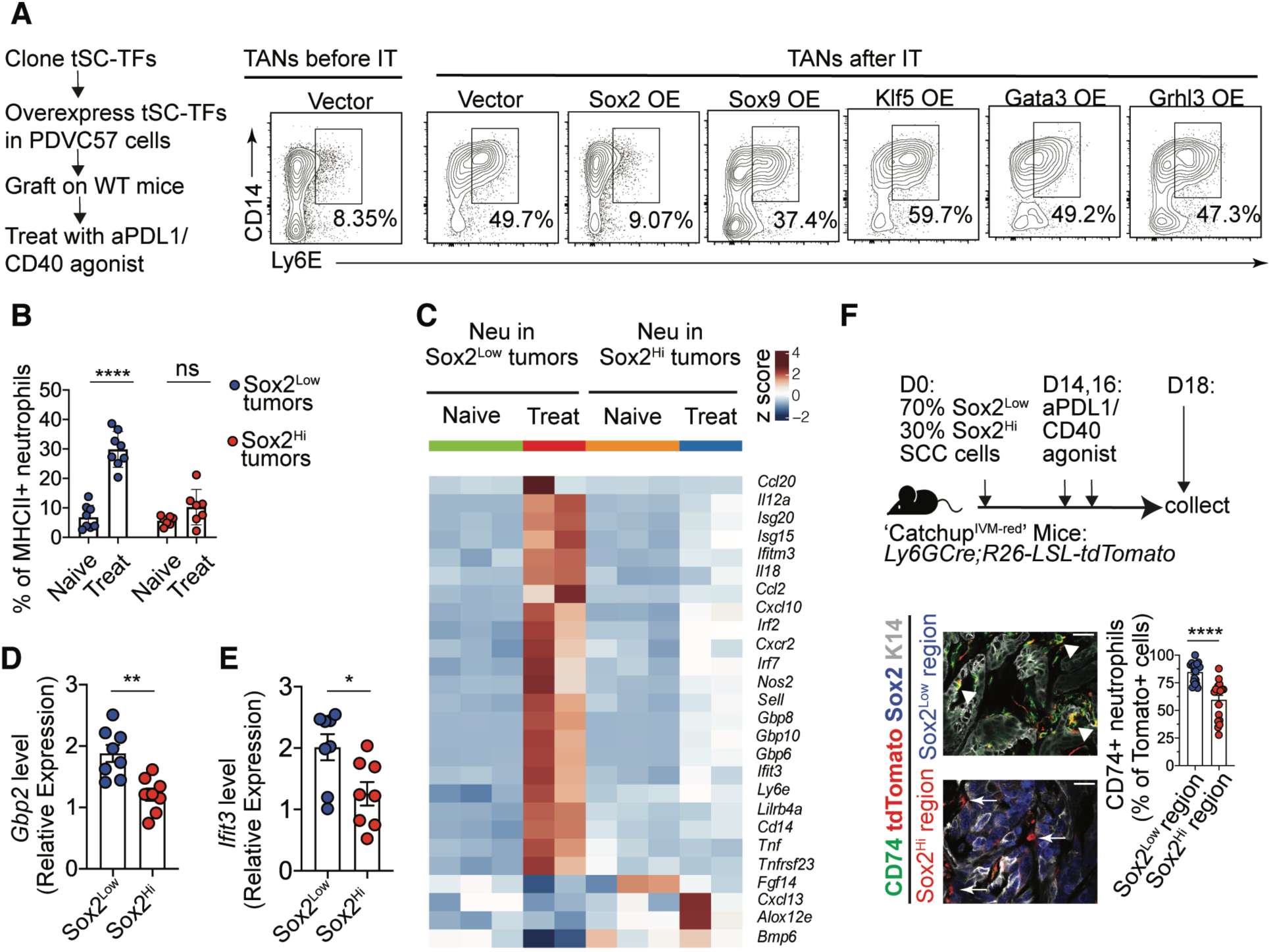
Sox2 amplification allows SCC cells to dampen the interferon-induced plasticity in TANs. **(A)** Experimental scheme and representative flow cytometry plots quantifying upregulation of CD14 and Ly6E in TANs isolated from tumors formed by SCC cells individually amplifying various TIC-specific TFs. **(B)** Flow cytometry quantification of the MHCII expression in TANs from SCC tumor cells with or without amplifying Sox2, before and after anti-PDL1/CD40 agonist treatments. n=7 in each group. **(C)** Heatmap showing the differentially expressed genes in neutrophils isolated from SCC tumors with (Sox2^Hi^) or without (Sox2^Low^) amplifying Sox2, before (Naïve) and after (Treat) anti-PDL1/CD40 agonist treatments. (**D** and **E)**. Quantitative PCR measuring the *Gbp2* (D) and *Ifit3* (E) expression in neutrophils isolated from naive Sox2^Low^ or Sox2^Hi^ tumors following *in vitro* IFNα/IFNγ treatments for 4 hr. n=8 in each group. **(F)**. Experimental scheme, representative IF images and quantification of CD74 expression in TANs enriched in Sox2 positive or low regions when Sox2^Low^ and Sox2^Hi^ cells were mixed at 7:3 ratio for grafting on neutrophil reporter mice. Scale bars: 50μm. Graphs show representative results from one the three repeats for each experiment and are presented as mean ± SEM. *p < 0.05; **p < 0.01; ****p < 0.0001; ns, non-significant.

Sox2 is a TF that plays a key role in embryonic stem cells and becomes reactivated in many epithelial cancers (*39–41*). In SCCs, Sox2 was demonstrated to be specifically enriched in TICs and was responsible for initiating and maintaining SCCs (*42*). Sox2 was expressed at a particularly high level in TGFβ-responsive TICs at the tumor-stroma interface (*37*). Thus, this intriguing finding from the reverse genetic screening prompted us to further test whether Sox2 is responsible for endowing TICs with the capacity to modulate neutrophils and suppress their plasticity. To address this question, we grafted either parental Sox2^Low^ control or Sox2^Hi^ SCC cells on the ‘Catchup^IVM-red^’ mice. Once the SCC tumors were established, we treated the mice bearing these two groups of tumors with anti-PDL1/CD40 agonist. We then isolated Tomato+ neutrophils from both naïve or treated Sox2^Low^ or Sox2^Hi^ SCC tumors and subjected them to bulk RNA-seq. Differential Gene Expression (DGE) showed that many anti-tumor immunity-related genes, including interferon responses (both type I and II) or antigen presentation pathways were strongly induced in neutrophils in parental Sox2^Low^ SCC tumors after immunotherapy (Fig. 4C). In contrast, the neutrophils found in the Sox2^Hi^ tumors were paralyzed and non-responsive and could not activate these anti-tumor functions (Fig. 4C). It was still possible that these blunted responses in neutrophils resulted from a lack of robust T cell responses in Sox2^Hi^ tumors, and hence a lack of sufficient interferons production in the TME to reprogram neutrophils, rather than a result of reduced plasticity. Although our RNAScope-based imaging analysis implied this was not the case (Fig. 3D), we sought to design additional experiment and test this possibility. To achieve this goal, we isolated TANs from untreated naïve Sox2^Low^ control or Sox2^Hi^ SCC tumors, and then treated them with interferons (IFNα + IFNγ) *in vitro*. Whereas the neutrophils from Sox2^Low^ tumors effectively activated interferon stimulated genes, the interferon responsive potential of TANs isolated from Sox2^Hi^ SCCs was significantly blunted (Fig. 4D,E). Importantly, when we isolated neutrophils from bone marrow or spleen from the same mice bearing the Sox2^Hi^ SCCs, their ability to respond to interferons appeared to be comparable to the neutrophils from the mice bearing Sox2^Low^ tumors (fig. S3A,B). Based on these results, we next designed a co-transplantation assay. To mimic the representation of Sox2+ stem cells in spontaneous tumors, we mixed Sox2^Hi^ SCC cells with parental Sox2^Low^ cells at a 3:7 ratio and then co-grafted the mixed SCC cells on the Catchup^IVM-red^’ mice, followed by anti-PDL1/CD40 agonist treatment and microscopy imaging. We found that only the neutrophils near the Sox2^Low^ cancer cells could effectively activate the class II antigen presentation machinery, as depicted by CD74 expression (Fig. 4F), whereas the Sox2^Hi^ cells could effectively block their surrounding TANs from becoming immune stimulatory (Fig. 4F).

Collectively, these key findings suggested that Sox2 amplification in SCC cells can effectively imprint neutrophils within their surroundings, limiting the neutrophils’ plasticity by preventing them from responding to interferons. Importantly, these effects of Sox2 are well conserved in different epithelial cancer types. When we amplified Sox2 in head and neck SCC cells followed by the anti-PDL1/CD40 agonist treatment, Sox2^Hi^ cancer cells could also efficiently block neutrophil responses, causing dampened T cell immunity (fig. S3C-H).

### Sox2^Hi^ TICs shape neutrophil cell states, block neutrophil-T cell interactions, exclude immune infiltration, and drive cancer relapse following immunotherapy

Our initial functional assays using grafted SCC cells expressing different levels of Sox2 provided solid evidence supporting its critical roles in facilitating cancer cells to modulate TANs. However, in autochthonous SCCs, Sox2 is only amplified in TICs (*42, 43*). Thus, it was imperative to further explore how Sox2^Hi^ TICs regulate neutrophils in a more physiologically relevant cancer model. To achieve this goal, we first generated *Krt14CreER; Sox2 ^flox/flox^* mice and used DMBA/TPA to induce autochthonous SCCs (Fig. 5A). When the tumors formed and progressed to SCCs, we injected tamoxifen to specifically ablate *Sox2* in basal skin epithelium where TICs reside (*Sox2* cKO), followed by treatments with anti-PDL1/CD40 agonist. To comprehensively investigate how the loss of Sox2 in TICs impact the TANs in SCCs, we subjected total CD45+ cells isolated from both naïve and treated *Sox2* cKO tumors to scRNA-seq. Strikingly, compared to the WT tumors in which a subset of less-plastic and interferon non-responsive TANs (T4 cluster) was preserved after immunotherapy treatment (Fig. 1C-F), the neutrophils infiltrating *Sox2* cKO tumors completely lost their heterogeneity (Fig. 5B). Additionally, almost all the neutrophils in the *Sox2* cKO tumors became interferon responsive with similar signatures of neutrophils of the T1 cluster in treated WT tumors (Fig. 5B). Notably, compared to the same T1 cluster in WT tumors, the interferon responses, immune co-stimulation, and class II MHC antigen presentation signatures were further enhanced in TANs when Sox2 was deficient in TICs (Fig. 5C).

**Fig. 5.**
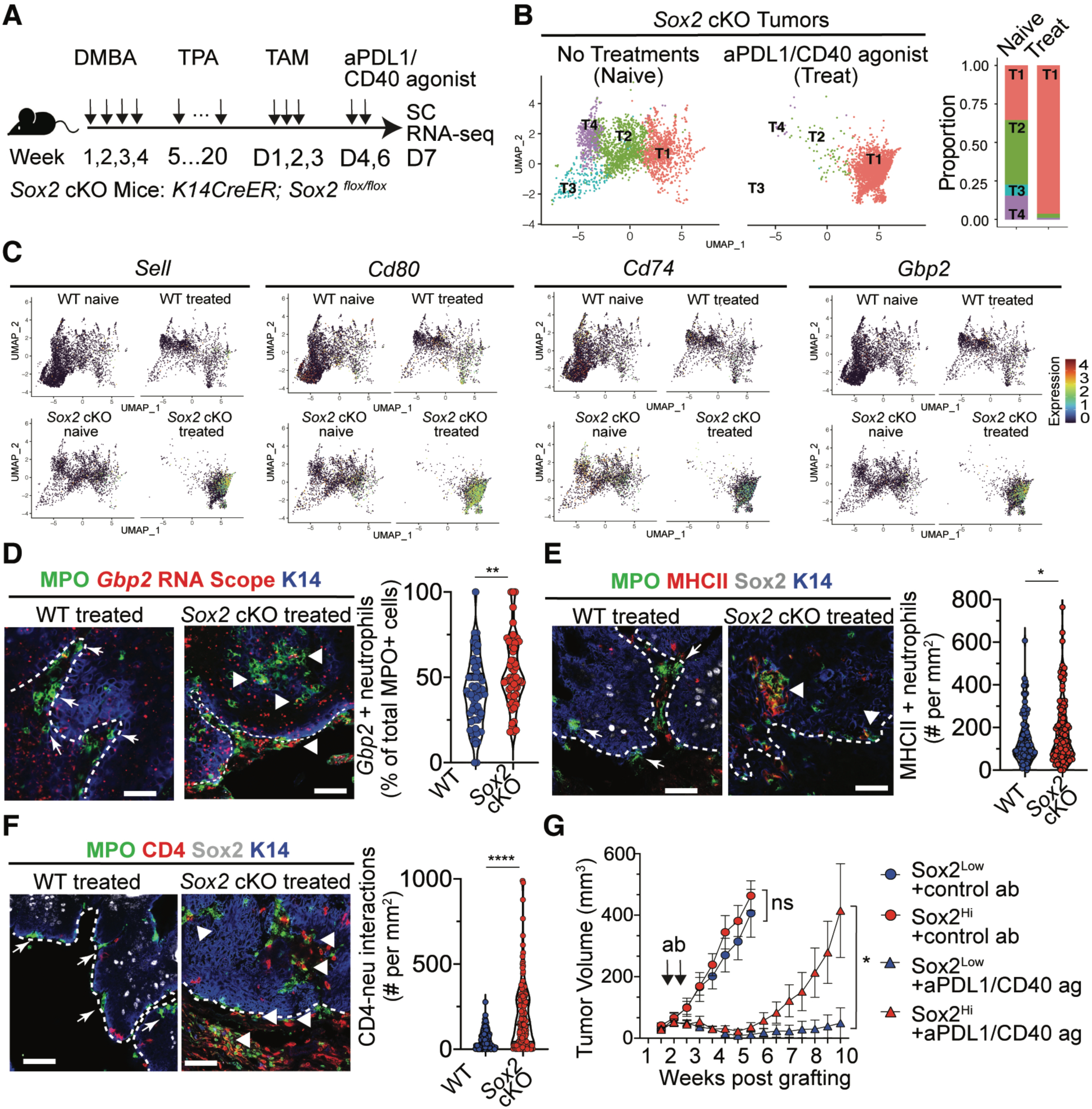
Sox2 is critical for TICs to shape neutrophil cell states, block neutrophil-T cell interactions, exclude T cells and drive cancer relapse. **(A)** Schematics showing the model for ablating Sox2 in TICs. **(B)** UMAP and stacked bar chart showing the changes in the composition of TAN subpopulations induced by immunotherapy when Sox2 is ablated in TICs. **(C)** UMAP showing expression of various immune stimulatory genes expressed in different TAN clusters in WT or *Sox2* cKO SCCs. **(D** to **F)**. Representative IF images and quantification of *Gbp2* expression (**D**), MHCII expression (**E**) and interactions with CD4 T cells (**F**) of MPO+ neutrophils in WT or *Sox2* cKO SCC tumors after anti-PDL1/CD40 agonist treatments. Tumor boarder was labelled with white dotted line. Scale bars: 50μm. **(G)** Growth of Sox2^Hi^ or Sox2^Low^ SCC cells, with or without anti-PDL1/CD40 agonist treatments. n=8 in each group. Representative results from one of the three repeats were presented. For each time point, the tumor sizes were shown as mean ± SEM. *p < 0.05; **p < 0.01; ****p < 0.0001; ns, non-significant.

By comparing the neutrophil landscape between Sox2 sufficient and deficient SCC tumors, it was clear that the Sox2-activated pathway could allow TICs to shape the cell states of TANs by modulating their plasticity. We next sought to determine the functional consequence of this stem cell-neutrophil crosstalk during immunotherapy treatments. Both spatial transcriptomic profiling and RNAScope-based imaging analysis have shown that most interferon responsive neutrophils were concentrated in the stroma in WT tumors (Fig. 3). When we examined the treated *Sox2* cKO tumors, we found that a large number of *Gbp2*+ neutrophils infiltrated the Sox2-deficient tumors (Fig. 5D). These neutrophils infiltrating the tumor mass also activated MHCII, indicating their potentials to prime or enhance CD4 T cells responses (Fig. 5E). As a result, we detected dynamic and intimate interactions between neutrophils and T cells in both the stroma, and within the tumor mass (Fig. 5F). Importantly, whereas most CD4 T cells were excluded from infiltrating the TIC-enriched regions in WT tumors, we can now detect strong T cell infiltration into the Sox2 cKO tumors (Fig. 5F).

Since previous studies have demonstrated that the TICs could survive immunotherapy even when effective treatments rapidly eliminated majority of differentiated tumor populations (*9*), we designed experiments to test whether Sox2 mediated the enhanced survival of TICs during immunotherapy treatments. For this purpose, we grafted either Sox2^Low^ or Sox2^Hi^ SCC cells on WT mice followed by either control antibody or anti-PDL1/CD40 agonist treatments. Whereas Sox2^Low^ and Sox2^Hi^ SCC tumors grew similarly in mice treated with control antibodies, and the growth of both SCC tumors could be initially reduced by the immunotherapy treatments, but only Sox2^Hi^ tumors quickly relapsed (Fig. 5G).

Taken together, we provided evidence that Sox2 amplification enables TICs to suppress the interferon-induced plasticity in TANs at the tumor-stroma interface. This immune modulatory activity of TICs is crucial during immunotherapy, as it allows TICs to manipulate a subset of TANs in their vicinity and block their response to treatments. This crosstalk preserves the immune suppressive state of these TANs and prevents them from priming tumor-infiltrating T cells. Altogether, this crosstalk allows TIC-modulated TANs to form a micro-niche that offers reciprocal protections for TICs against T cell-mediated attacks.

### Sox2 activates Fads1 in TICs to block the interferon responses in TANs

Built on these findings, which supported a previously unrecognized immune modulatory roles of Sox2 in cancers, we explored the underlying mechanisms activated by Sox2 in TICs that was responsible for modulating neutrophils. We speculated that the Sox2 amplification must induce the secretion of certain factors from TICs that can disrupt the interferon responsiveness in neutrophils. To identify such unknown factors, we grafted and isolated Sox2^−/-^ and Sox2^Hi^ SCC cells, and then subjected these cells for bulk RNA-seq. We were particularly intrigued by the enhanced expression of *Fads1* in Sox2^Hi^ cells (Fig. 6A). We further confirmed the direct regulation of *Fads1* gene expression by Sox2 using CUT & RUN-sequencing which identified the direct binding of Sox2 at the promoter region of *Fads1* gene locus (Fig. 6B). Fads1 is a Δ5 desaturase that can catalyze linoleic and linolenic acids metabolism to produce polyunsaturated fatty acids (PUFA), such as arachidonic acid (AA) (*44*). AA is a highly bioactive molecule that could have profound impacts on immune cell physiology (*45, 46*). To investigate the functional significance of Fads1 in Sox2^Hi^ SCC cells’ modulation of TANs, we first performed CRISPR gene editing and generated *Fads1*^−/-^ PDVC57 cells, and then amplified *Sox2* in these cells. After engraftment and immunotherapy treatment, we analyzed neutrophil phenotypes. Interestingly, silencing *Fads1* in Sox2^Hi^ SCC tumors reactivated the immunotherapy-induced reprogramming in TANs (Fig. 6C, D). Even though Sox2 was still overexpressed in the same cancer cells, the TANs infiltrating these tumors could now upregulate Ly6E or class II MHC upon receiving anti-PDL1/CD40 agonist treatments (Fig. 6C, D). Importantly, when *Fads1* was silenced, the neutrophils from these Sox2^Hi^ SCC tumors could now respond to interferons when isolated and treated with interferons *in vitro* (Fig. 6E).

**Fig. 6.**
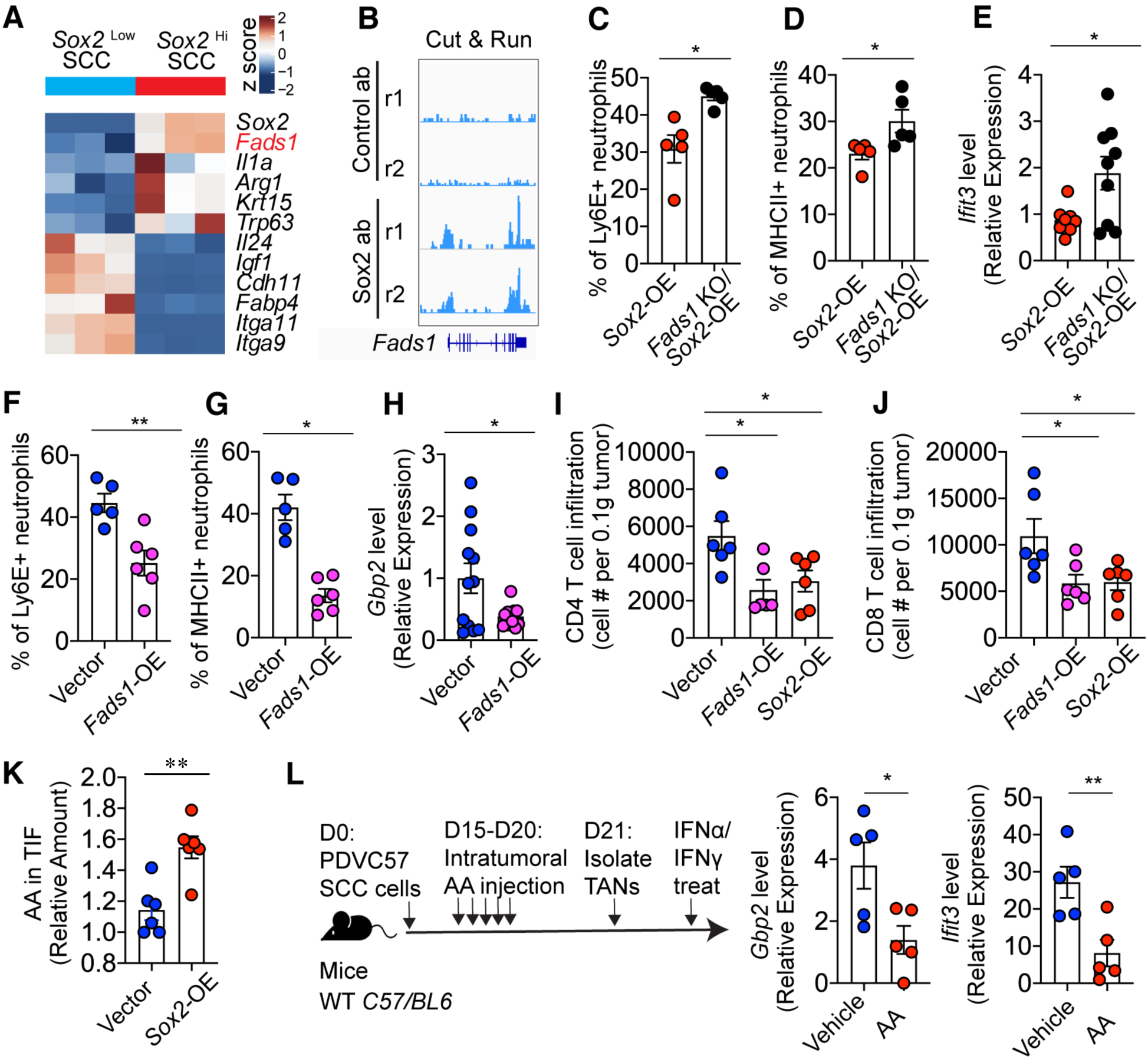
Sox2 activates Fads1 to produce arachidonic acid to block the interferon-induced reprogramming in TANs. **(A)** Heatmap showing the differentially expressed genes in Sox2^Hi^ SCC cells compared to Sox2 KO cells. **(B)** IGV image showing the Sox2 or control antibody CUT & RUN. Note the positive binding of Sox2 over the *Fads1* gene locus. **(C** to **D)**. Flow cytometry quantification of the Ly6E (**C**) and MHCII (**D**) expression in TANs in immunotherapy-treated SCC tumors formed by Sox2^Hi^ cells with or without silencing *Fads1*. n=5 in each group. **(E)** Quantitative PCR measuring *Ifit3* expression in TANs isolated from naïve SCC tumors formed by Sox2^Hi^ cells with or without silencing *Fads1* following treatment with IFNα/IFNγ *in vitro* for 4 hr. n=9 in each group**. (F** to **G)** Flow cytometry quantification of the Ly6E (**F**) and MHCII (**G**) expression in TANs in immunotherapy-treated SCC tumors formed by SCC cells with or without amplifying *Fads1*. n=5 in each group. **(H)** Quantitative PCR measuring *Gbp2* expression in TANs isolated from naïve SCC tumors with or without amplifying *Fads1* following treatment with IFNα/IFNγ *in vitro* for 4 hr. n=12 in each group. **(I** to **J)** Flow cytometry quantification of CD4 T cell infiltration (I) and CD8 T cell infiltration (J) in immunotherapy-treated SCC tumor cells formed by SCC cells with or without amplifying *Fads1*. n=6 in each group. **(K)** Quantitative mass spectrometry measurement of AA in TIF extracted from naïve SCC tumors formed by Sox2^Low^ or Sox2^Hi^ tumors. **(L)** Experimental scheme and quantitative PCR measuring the *Gbp2* and *Ifit3* expression in TANs isolated from SCC tumors with or without intratumoral injection of AA. Isolated neutrophils were treated with IFNα/IFNγ *in vitro* for 4 hrs to measure their interferon responsive capacity. n=5 in each group. Bar graphs show representative results from one of the two or three repeats for each experiment and are presented as mean ± SEM. *p < 0.05; **p < 0.01.

In parallel, we amplified *Fads1* in Sox2^Low^ PDVC57 SCC cells and treated mice bearing these tumors with anti-PDL1/CD40 agonists. Interestingly, *Fads1* amplification in SCC cells alone was sufficient to enable cancer cells to manipulate TANs and block them from responding to interferons. For example, similar to the TANs in Sox2^Hi^ tumors, the TANs in the Fads1^Hi^ tumors also maintained their Ly6E ^low^ immune suppressive phenotypes (Fig. 6F), and these TANs could not activate class II antigen presentation (Fig. 6G). In addition, we isolated neutrophils from naive Fads1^Hi^ tumors and treated them with interferons *in vitro*. Compared to the neutrophils from vector control tumors, the neutrophils isolated from the Fads1^Hi^ tumors could not efficiently induce interferon-stimulated genes (Fig. 6H). As a result, similar to the T cells infiltrating the Sox2^Hi^ tumors, T cell infiltrations were also significantly blunted in the Fads1^Hi^ tumors (Fig. 6I, J). Therefore, we conclude that Fads1 is the key downstream effector activated by Sox2 in TICs to modulate the interferon responsive potentials of TANs.

### Sox2^Hi^ TICs produce AA to block interferon response in TANs

The significance of the TIC-specific Sox2-Fads1 axis in modulating the TANs’ plasticity prompted us to further investigate whether AA plays a key role in mediating the suppression of interferon responsiveness in TANs. We first investigated whether Sox2 amplification allowed SCC cells to produce more AA. For this purpose, we focused on the tumor interstitial fluid (TIF). TIF perfuses through the tumor, facilitating tumor cell exchange of nutrients, metabolites, cytokines, and signaling molecules (*47*). Notably, the composition of metabolites and signaling factors in TIF faithfully reflect the molecular components present in the TME. As we expected, when we isolated TIF from SCC tumors formed by either Sox2^low^ parental control or Sox2^Hi^ cells, quantitative mass spectrometry analysis shows AA is significantly enriched in the TIF from Sox2^Hi^ tumors (Fig. 6K). AA is an essential PUFA that has versatile bioreactivity and can modulate various physiological conditions (*48*). However, to our knowledge whether AA can block cellular responses to interferons has not been reported before. To explore this new function of AA, we intratumorally injected AA into the SCCs formed by Sox2^Low^ PDVC57 cells, isolated TANs from vehicle or AA treated tumors, and treated these neutrophils *in vitro* with interferons (Fig. 6L). Importantly, this key experiment confirmed that presence of high amount of AA in the TME can effectively blunt the interferon responsive capacity of TANs.

Cumulatively, these results place the TICs at the center stage of shaping the plasticity and heterogeneity of TANs. Our new finding not only provide compelling evidence supporting the critical roles of TICs in modulating TANs during immunotherapy treatment, but also identified a new stem cell-specific pathway that is orchestrated by Sox2, which activates Fads1 to increase the local production of AA. Uptake of the excessive amount of AA by TANs can disrupt the interferon responsive potentials in TANs. This intimate crosstalk initiated by TICs prevented a subset of TANs near TICs from gaining plasticity and being reprogrammed by immunotherapy-induced interferons, thus, preserved their immune suppressive activity. Thus, TICs can manipulate this subset of TANs to form a protective micro-niche at the tumor stroma interface that could shield TICs from infiltrating T cells and eventually drive tumor relapses.

## Discussion

Stem cells, as long-lived cells governing the regeneration of various tissues, have long been considered as a group of vulnerable cells that must be constantly protected within an immune-privileged niche (*14, 49*). Our study has now challenged this paradigm and found that instead of passively receiving protections, stem cells are wired to modulate the cell states and differentiation trajectory of immune cells and actively sculpt their own dynamic niche within a highly inflammatory environment. Importantly, this stem cell feature can be hijacked by TICs to shape the suppressive tumor microenvironment. Supporting this idea, we have identified a new crosstalk initiated by TICs to manipulate TANs, allowing the neutrophils to build a protective-niche during immunotherapy. TICs comprise a vital cellular reservoir for repopulating the tumors when effective therapies remove the bulk of the tumor mass (*7, 50*). Thus, TICs must be equipped with special molecular programs to orchestrate therapy resistance and cancer relapse. Unlike chemo- or radiation therapies, which only generate transient effects during the active treatments, effective immunotherapies are supposed to elicit long-lasting changes in the TME (*51*), activate stable tissue residency programs (*52*), and establish long-term immune memory within infiltrating immune compartments (*53*). It remains unclear how TICs can overcome these enduring effects from successful immunotherapy and eventually give rise to the relapsed tumors after the immunity is reinvigorated. In this study, we have unveiled how TICs could achieve this capability. We found that TICs can fine-tune the plasticity of a small subset of neutrophils that are recruited to their surroundings at the tumor-stroma interface, therefore, restraining these neutrophils from being reprogrammed by immunotherapy. This key modulation maintains the suppressive cell state in this subset of neutrophils and allows these neutrophils to form a micro-niche which, in turn, protects the TIC from being attacked by the robust anti-tumor T cell response evoked during the effective immunotherapy treatments.

Delving into the mechanisms driving this TIC-shaped protective niche, we have identified a new molecular circuit orchestrated by Sox2 for TICs to modulate TANs. Our study finds that Sox2 can activate Fads1 to produce arachidonic acid, which limits the reprogramming effects of interferons when taken-up by TANs. As the important precursor for producing prostaglandin and leukotriene (*45*), the immune modulatory actions of AA have been recognized(*48*). In particular, it has been shown that elevated expression of Fatty Acid Transport Protein 2 (FATP2) can facilitate TAN to import more AAs, allowing them to upregulate *Ptgs2* expression (*30*) and become immune suppressive in the TME. This study first identified TICs as the previously unknown source of enforcing a subset of TANs to take up more AA than the others. Secondly, this study identified a new role of AA in antagonizing the interferon responsive potentials and modulating the cellular plasticity of neutrophils. Although not the focus of this study, future research is needed to elucidate the molecular mechanisms by which AA disrupts interferon signal transduction in various immune cells.

Furthermore, our study has important implications for understanding the biology of TANs. The versatile roles of TANs during successful immunotherapy treatment have only been recently appreciated. How TANs switch phenotypes from being predominantly immune suppressive to supporting anti-tumor immunity during the immunotherapy treatment is still unknown. Although lacking definitive evidence to support, these anti-tumor activities were attributed to the newly recruited neutrophils attracted by effective immunotherapies (*23*). This notion was partially supported by recent studies proposing a model that neutrophils must undergo deterministic and irreversible epigenetic changes once they enter the TME, thereby lacking high degree of plasticity

(*16*). However, our *in vivo* lineage tracing and *in vitro* cytokine treatments support a different scenario, namely that TANs that are already present in the tumor stroma before treatments can still be reprogrammed by the immunotherapy-induced interferons, become more plastic, and be converted into anti-tumor neutrophils. Interestingly, it appears that the synergistic effect of both type I and II interferons is required to reprogram TANs (*22*). Future research is required to untangle the distinct and combined effects of IFNα/β and IFNγ on TANs. Comprehensive epigenetic profiling is also required to understand the impacts of interferons on the plasticity and cell states of TANs.

In closing, we have identified an intimate crosstalk between a subset of tumor-associated neutrophils and tumor-initiating stem cells at the tumor stroma interface, which can prevent these neutrophils from being reprogrammed by immunotherapy. As a result, they protect TICs from being attacked by infiltrating T cells. Thus, this study provides new insights for designing new strategies that can disrupt the dialogue between neutrophils and TICs, making TICs vulnerable to anti-tumor immunity and hence preventing cancer relapse from immunotherapy treatments.

## Acknowledgments

We thank J. Stanisavic in the Miao lab for assistance; C. Ciszewski at the Human Disease & Immune Discovery Core Facility at the UChicago for conducting FACS sorting; P. Faber (Genomics Core at the UChicago) for sequencing and raw data processing, H. Shah from The Metabolomics Platform, and Animal Resources Center and Special Service at UChicago for maintaining mouse colonies.

## Funding

This study was supported by Y.M.’s Start-up fund and seed grants from The University of Chicago, grants to Y. M. from NIH (R00CA237859, R01CA285786), V Foundation, American Association for Cancer Research, and The Cancer Research Foundation.

## Author contributions

Y.M., W.G. and J.L., conceptualized the study, designed the experiments, interpreted the data, and wrote the manuscript. W.G. and J.L., performed most experiments and analyzed data with the assistance from H.X., L.D., G.J., N.B., L.B., O.A., I.E., R.A., A.N., B.D.. G.M. generated the Catchup mice, B.I. and H.A. generated Ly6GiCre mice. Competing interests: Authors declare that they have no competing interests.

## Data and materials availability

Raw and analyzed data is available at NCBI and the accession number for the single cell RNA sequencing, bulk RNA sequencing, Spatial transcriptomics profiling, CUT & RUN-seq data reported in this paper are NCBI GEO: GEO: GSE278435, GEO: GSE278436, GEO: GSE278430, GEO: GSE278429, GEO: GSE278426.

## Materials and Methods

### Animals

K14CreER mice were generated by Dr. Elaine Fuchs lab and backcrossed to C57/BL6J background for ten generations ^57^. *C57BL/6-Ly6g(tm2621(CretdTomato)Arte);R26-LSL-tdTomato* (Catchup ^IVM-red^) mice were generated by Dr. Matthias Gunzer’s lab at University Hospital Essen (PLI). *C57BL/6-iLy6GtdTom* mice were generated by Dr. Andrés Hidalgo’s lab at Yale University. These mice have been described before ^37^. Wild-type C57BL/6J, *IfngR ^flox/flox^ (C57BL/6N-Ifngr1tm1.1Rds/J), Sox2 ^flox/flox^ (B6.Sox2tm1.1Lan/J), S100a8Cre (B6.Cg-Tg(S100A8-cre,-EGFP)1Ilw/J), iDTR (C57BL/6-Gt(ROSA)26Sortm1(HBEGF)Awai/J)* mice, were obtained from The Jackson Laboratory. Neutrophils were depleted in *S100A8Cre;iDTR* mice by intraperitoneally (i.p.) injection of 0.25 μg diphtheria toxin (DT, Sigma-Aldrich) every 2 days for three doses. To treat mice with tamoxifen to induce Sox2 conditional knockout, a daily i.p. injection of 100 μg for three consecutive days was performed.

All mice were maintained in an Association for Assessment and Accreditation of Laboratory Animal Care (AAALAC)-an accredited animal facility. Procedures were performed using IACUC-approved protocols. All mice were maintained and bred under specific-pathogen free conditions at the University of Chicago. All the procedures are in accordance with the Guide of the Care and Use of Laboratory Animals.

### Cell lines

Mouse skin SCC line PDVC57 for tumor grafting and HEK293T cells for packaging lentivirus were cultured in Dulbecco’s modi’ed eagle medium (DMEM) in 10% FBS, 100 units/mL streptomycin and 100 mg/mL penicillin with 2mM glutamine. Mouse oral SCC line MOC1 was cultured in Iscove’s Modified Dulbecco’s Medium (IMDM)/F12 (2:1) media with 5% FBS, 100 units/mL streptomycin and 100 mg/mL penicillin, 5μg/mL insulin, 40μg/mL hydrocortisone, 5μg/mL EGF.

### Tumor formation and treatment

A two-stage chemical carcinogenesis protocol was used to induce skin SCCs in mice. The dorsal skin of various mouse strains was shaved and topically treated with 200 µL of 200nM 7,12-dimethylbenz[a]anthracene (DMBA) once a week for 4 weeks to initiate tumorigenesis. Four weeks later, tumor promotion was induced by weekly (twice a week) applications of 200 µL of 35µM 12-O-tetradecanoylphorbol-13-acetate (TPA) for 20 weeks. For tumor transplantation, 5×105 PDVC57 mouse skin SCC cells or 2×106 MOC1 mouse oral SCC cells were mixed with Cultrex Basement Membrane Extract, PathClear, Type 3 matrigels (Bio-Techne) and injected subcutaneously. Tumors were allowed to grow for two weeks prior to immunotherapy.

For immunotherapy treatment on both DMBA/TPA generated spontaneous tumors or grafted SCCs, tumor bearing mice were treated with a combination of PD-L1 blocking antibody (Clone 10F.9G2, 100 µg/mouse, BioXCell) and CD40 agonist antibody (Clone FGK4.5, 100 µg/mouse, BioXCell) intraperitoneal for a total of 2 doses every other day. Tumors were collected 2 days after the last treatment for downstream experiments. For intratumoral treatment with arachidonic acid, tumors were allowed to grow for three weeks prior to treatment. AA was mixed in 5% DMSO/PBS solution and 1.5 mg AA was injected into each tumor once a day for 5 consecutive days^58^.

### Cell sorting and flow cytometry

To sort immune cells from DMBA/TPA induced tumors, tumors were dissected at designated time points, minced and digested with 0.25% type 4 collagenase (GIBCO) in RPMI-1640 (GIBCO) for 20 minutes followed by 0.25% Trypsin treatment for 10 minutes at 37 °C. Digested tissues were then passed through a 70 µm cell strainer, subjected to ACK lysis for 1 minute to lyse the red blood cells and resuspended in PBS. Single cells suspension was incubated with Fc Block TruStain FcX (Clone 93, Biolegend) in PBS with 5% normal rat serum and 5% normal mouse serum, and then stained with a cocktail Ab at predetermined concentration in staining buffer (PBS with 5% FBS and 1% HEPES). DAPI was used to exclude dead cells. The sorting was performed on BD Symphony S6 Cell Sorter.

To pro’le immune population and measure tumor-in’ltrating T cell activity in grafted tumors, the tumors were minced and digested in 0.25% type 4 collagenase (GIBCO) in RPMI-1640 for 60 minutes at 37 °C with shaking. Digested tissues were then passed through a 70 µm cell strainer and red blood cells were lysed using ACK lysis buffer for 1 min. After preparation of single cell suspension, cells were then either treated with Cell Stimulation Cocktail (plus protein transport inhibitors) (Thermo Fisher) for 4 hours at 37 °C or directly stained Zombie Aqua (Biolegend) to exclude dead cells, Fc blocked with TruStain FcX, and then with a cocktail of Abs for surface antigens at pre-determined concentrations. The stained cells were run on LSR-Fortessa analyzer (BD Biosciences) and analyzed with FlowJo software.

### Neutrophil isolation

The neutrophil isolation methods were adopted from previous published protocols^59^. Briefly, to isolate neutrophils from grafted tumors, the tumors were minced and digested in 0.25% type 4 collagenase (GIBCO) in RPMI-1640 for 60 minutes at 37 °C with shaking. Digested tissues were then passed through a 70 µm cell strainer and red blood cells were lysed using ACK lysis buffer for 1 min. After preparation of single cell suspension, cells were resuspended in 8 ml of 40% Percoll and overlayed onto 3 ml of 70% Percoll in a 15 ml conical centrifuge tube. After centrifugation for 30 minutes at 900 × g without brake at room temperature, cells from the 70%/40% Percoll interface, were harvested, washed twice with FACS buffer and resuspend in FACS buffer. Cells were blocked with TruStain FcX for 10 min, then incubated with 20 µL biotin-anti-Ly6G antibody for 15 min followed by incubation with 20 µL magnetic strep-beads for 15 min. Samples were washed 4 times with FACS buffer and resuspended in RPMI-1640 for *in* vitro interferon treatments.

To isolate neutrophils from mouse bone marrow or spleen, femurs were harvested, and both ends were cut using sterile scissors. The bone marrow was flushed from the femurs using a syringe filled with RPMI-1640 medium and passed through a 40 µm cell strainer into sterile tubes. The spleen was harvested, mechanically dissociated, passed through a 40 µm cell strainer, and collected into sterile tubes. The collected cells were treated with ACK lysis buffer for 1 minute to lyse red blood cells. Following lysis, cells were blocked and subjected to the same isolation procedure described for tumor samples.

### Neutrophil culture and treatments

Freshly isolated neutrophils were cultured in RPMI-1640 with 10% FBS, 100 units/mL streptomycin and 100 mg/mL penicillin, 55 µM beta-mercaptoethanol (GIBCO), 50U/mL GM-CSF (Biolegend). For *in vitro* IFN pathway activation, 5 µg /mL of IFN-α and 5 µg/mL of IFN-γ were added to the culture media and incubated for 4 hours prior to harvest cells for further analysis.

### Cloning and transducing cells

Mouse Sox2 and Fads1 full open reading frames (ORFs) were amplified from PDVC57 cDNA through PCR, then cloned into lentiviral vectors using NEBuilder® HiFi DNA Assembly kit (New England Biolabs) following the manufacturer’s instructions.

Virus was produced by co-transfecting the viral plasmids, psPAX2, and pMD2.G plasmids using Lipofectamine 3000 (Thermo Fisher), according to the manufacturer’s instructions. The medium was replaced 4 hours post-transfection. Viral supernatants were collected 48 hours post-transfection, filtered through a 0.45 µm filter.

For transduction, target cells were seeded in 6-well plates. Viral supernatant was added with 10 µg/mL polybrene. Plates were centrifuged at 1,100 × g for 30 minutes at 37°C. After centrifugation, the viral medium was replaced with fresh medium, and transduced cells were cultured for 48 hours before further analysis or selection.

### TIF extraction

Immediately after tumor dissection, tumors were cut into small pieces and then place on 40um cell strainer in a 50 mL conical tube and centrifuged at 300g for 10 min at 4°C. The AA in TIF isolated from different tumors were quantified by mass spectrometry.

### RNA isolation and sequencing library preparation

Total RNAs from FACS-sorted cells were extracted using Quick-RNA Microprep Kit (Zymo Research) following the manufacturer protocol. RNA-seq libraries were prepared with the NEBNext Single Cell/Low input RNA library prep kit for Illumina (NEB) following the manufacturer protocol. The libraries were sequenced on the Illumina Novaseq platform.

### Single cell cDNA synthesis and sequencing library preparation

Tumors were collected 1 day after last dose of immunotherapy and total CD45 positive immune cells were sorted. Immediately after sorting, the sorted immune cells were fixed with Evercode Cell Fixation v2 (Parse Biosciences). the fixed cells were prepared using Evercode WT v2 kit (Parse Biosciences) according to manufacturer’s instructions. Input cell numbers and volumes were calculated using WT_100K_Sample_Loading_Table_V1.3.0 (Parse Biosciences). All the samples were processed together with Evercode WT v2 kit (Parse Biosciences) and 8 sub-libraries were generated. The quality of sub-libraries was checked by the Agilent Tape Station system (Agilent) and sequenced by Illumina NovaSeqX.

### CUT&RUN library preparation

CUT&RUN was performed using CUTANA™ ChIC/CUT&RUN Kit version 3 (EpiCypher) following the manufacture protocol. For each CUT&RUN reaction, 500,000 PDV-Sox2 cells were first cross-linked using 1% formaldehyde for 1 min at room temperature and then quenched by adding 2.5 M Glycine to a final concentration of 125 mM. Crosslinked cells were permeabilized, immobilized onto Concanavalin-A beads and incubated overnight (4 °C with gentle rocking) with either rabbit IgG or Sox2 antibody. DNA was extracted after chromatin digestion and release. Sequencing libraries were prepared with the NEBNext® Ultra™ II DNA Library Prep Kit for Illumina (NEB) according to the manufacture protocol.

### NanoString GeoMx Mouse Whole Genome Transcriptome Profiling

NanoString’s GeoMx Digital Spatial Profiler (DSP) platform was used to perform whole-genome transcriptome profiling on mouse tissue samples. Fresh frozen tumor sections were mounted onto slides, followed by immunofluorescent staining with specific antibodies targeting protein markers of interest to delineate distinct tissue regions. After staining, high-resolution digital images were acquired to visualize and identify regions of interest (ROIs).

ROIs were selected based on morphological and biological relevance. These regions were then subjected to UV light-based spatially resolved photocleavage to release oligonucleotide tags bound to the target mRNA transcripts. The released tags were collected were collected into 96-well plates. Library preparation was performed according to the NanoString GeoMx-NGS Readout Library Prep manual. Briefly, the released tags were dried and reconstituted in 10 µL nuclease free water, and 4 µL was used in a PCR reaction. NanoString barcoded primers were used to amplify the tags and add Illumina adaptor sequences and sample demultiplexing barcodes. PCR products were pooled in equal volumes and purified with two rounds of AMPure XP beads (Beckman Coulter). Libraries were sequenced on an Illumina NextSeq 550 platform.

### Immunofluorescence and RNA scope staining

Tumors were fixed in 1% paraformaldehyde for 1 hour at 4°C and washed with cold PBS for three times. Followed by incubation in 30% sucrose at 4°C overnight, tumor tissues were embedded in OCT (Tissue Tek), frozen, and sectioned (10-16 um). Cryosections were permeabilized, blocked, and stained with the following primary antibodies: CD4 (rat, 1:100, Biolegend), Krt14 (chicken, 1:1000, Biolegend), SOX2 (rabbit, 1:400, Cell Signaling Technology), SOX2 (rat, 1:100, ThermoFisher Scientific), RFP (rat, 1:1000, Proteintech), MPO (goat, 1:40, R&D), CD74 (sheep, 1:20, R&D), I-A/I-E (rat, 1:200, Biolegend), or Cleaved caspase3 (rabbit, 1:500, Cell Signaling Technology). The samples were then stained with related secondary antibodies conjugated with Alexa Fluor 488, Rhodamine Red, or Alexa Fluor 647 (Jackson ImmunoResearch Laboratories) and imaged on Leica Stellaris 8 Laser Scanning Confocal microscope.

For the RNA scope staining, cyrosections were dehydrated, performed with hydrogen peroxide and Protease III, hybridized with AMPs and developed the HRP-C1 channel with the Gbp2 probe according to the RNAscope Multiplex Fluorescent Reagent Kit Assay (ACDbio). Immediately after RNA labeling, the cytosections were blocked and stained with the primary antibodies against Krt14 and MPO at 4°C overnight and then incubated with the related secondary antibodies. The images were collected by Leica Stellaris 8 Confocal microscope and analyzed using Fiji/ImageJ software.

### RNA puri’cation and qRT-PCR

The total RNA was puri’ed using Quick-RNA Microprep Kit (Zymo Research) in accordance with manufacturer’s instructions. For real-time qRT-PCR analysis of any target genes, equal amount of RNAs were reverse transcribed using Maxima First Strand cDNA Synthesis Kit (Thermo Fisher). cDNAs were then mixed with speci’c gene primes and PowerTrack™ SYBR Green Master Mix (Thermo Fisher) and the qRT-PCR was performed on the QuantStudio™ 3 Real-Time PCR System (Thermo Fisher).

### CRISPR mediated gene knockout

To generate gene knockouts, the lentiCRISPRv2 system was employed. Specific single guide RNAs (sgRNAs) targeting the gene of interest were designed using the CRISPick (https://portals.broadinstitute.org/gppx/crispick/public) and cloned into the lentiCRISPRv2 vector (Addgene #52961)^60^.

HEK293T cells were transfected with the lentiCRISPRv2 construct, along with packaging plasmids (psPAX2 and pMD2.G) to produce lentiviral particles. After 48 hours, viral supernatants were collected, filtered, and used to transduce target cells in the presence of 10 µg/mL polybrene to enhance infection efficiency.

Following transduction, cells were selected with puromycin (2 µg/mL) for 3days to enrich for successfully transduced cells. Single clones were generated by limiting dilution.

### Single-cell RNA-seq data analysis

The raw FASTQ ’les were processed by spipe.v1.1.1(Parse Biosciences) using mm10 as a reference. Count matrices were imported to Seurat (v.4.3.0).^61^ Cells with <50 detected genes or >10000 detected genes or > 7.5% mitochondrial genes were filtered-out from the dataset. We then used the standard Seurat pipeline by running NormalizeData(), FindVariableFeatures(selection.method =”vst”, nfeatures =4000), and ScaleData(). For the first round of clustering, Principal component analysis (PCA) was performed and the top 50 PCs with a resolution = 0.6 were applied. RunUMAP(dim=1:40, n.neighbours =30) was used to visualize the data. Cell types were annotated using SingleR (V1.7.1)^62^ package with ImmGen data as reference. Neutrophils were subset, and standard pipeline was applied to neutrophil population by running FindVariableFeatures(selection.method =”vst”, nfeatures =4000), and ScaleData(). For clustering the neutrophils, Principal component analysis (PCA) was re-performed and the top 40 PCs with a resolution = 0.2 were applied. For visualization, RunUMAP(dim=1:50, n.neighbours =30) was used.

For RNA velocity analysis, spliced and unspliced count matrices were first obtained from spipe output and exported as h5ad files for subsequent input of scVelo. Velocity analysis was then performed with scVelo (0.2.5)^63^ using the default stochastic model and velocity vectors were projected into the UMAP generated from previous Seurat pipeline.

For pesudobulk analysis, cells were randomly grouped into 3 pseudo replicates. Gene counts were summarized and DESeq2 R package (v1.30.0)^64^ in R (v4.1.1) was used for differential gene analysis.

### NanoString GeoMx data analysis

FASTQ files were first processed by the NanoString GeoMx NGS Pipeline v2.0. Briefly, raw reads were pre-processed by trimming low-quality bases and adapter sequences. Filtered reads were then stitched and aligned, followed by extraction of barcode and UMI sequences. Barcodes were matched to a reference set of known probe barcodes, allowing for a maximum of one mismatch. To remove duplicates, reads sharing the same barcode were collapsed based on unique molecular identifiers (UMIs).

Raw data from NanoString GeoMx NGS Pipeline were then imported to R using GeomxTools package to generate GeoMxSet object. The raw count data was first normalized with “q_norm” method. To facilitate data processing, GeoMxSet object was further coerced into Seurat object and standard Seurat pipe lines were applied for analysis.

### Bulk RNA-Seq Alignment and Differential Expression Analysis

Raw FASTQ files were first trimmed and filtered by cutadapt (v3.2)^65^. Estimated transcript counts for the mouse mm10 genome assembly (GENCODE vM24) were obtained using the pseudo-aligner Kallisto (v0.44.0)^66^. Gene level abundance was summarized and differential gene expression was performed using the DESeq2 R package (v1.30.0) in R (v4.1.1). Genes with Benjamini-Hochberg method adjusted P values < 0.05 were regarded as significantly differentially expressed.

### Gene Set Enrichment Analysis (GSEA)

GSEA was performed using clusterProfiler R package (v4.6.0, https://guangchuangyu.github.io/software/clusterProfiler)^67^. MSigDB (Molecular signature database, v7.4) gene set collections (http://www.gsea-msigdb.org/gsea/msigdb/index.jsp)^68^ was used in the analysis. Genes were ranked by the fold change value obtained from DESeq2. Enriched pathways with Benjamini-Hochberg method adjusted P values < 0.25 were considered to be significant.

### Statistics

Data are presented as mean ± SEM or mean ± 95% con’dence interval. P values were determined by using two-tailed Student’s t test, Mann-Whitney U Tests or ANOVA test. Statistical analyses were performed using Prism 9 (GraphPad). Experiments were performed in an open-label manner. Signi’cant difference between two groups were noted by asterisks (* p < 0.05; ** p < 0.01: *** p < 0.001).

---

**fig. S1.**
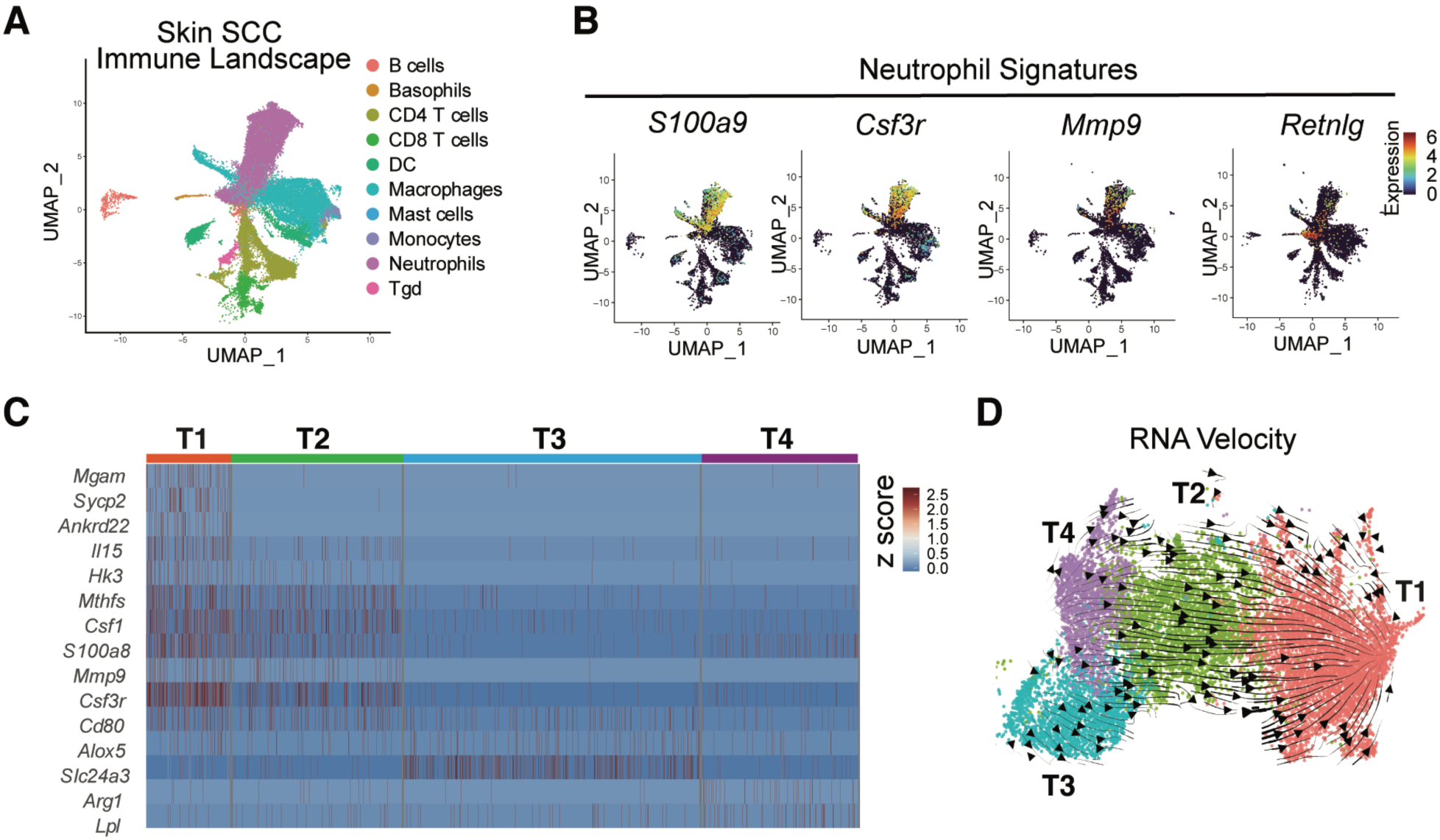
Single cell analysis of T cell and neutrophil responses in skin SCC. Related to Figure 1. **(A)** UMAP showing the immune cell types identified by scRNA-seq in skin SCCs. **(B)** UMAP showing the expression of signature genes used for identifying the neutrophils in skin SCCs. **(C)** Heatmap showing the expression of signature genes enriched in different clusters of TANs**. (D)** RNA velocity analysis of potential developmental trajectory and relationship among various neutrophil clusters in skin SCCs.

**Fig. S2.**
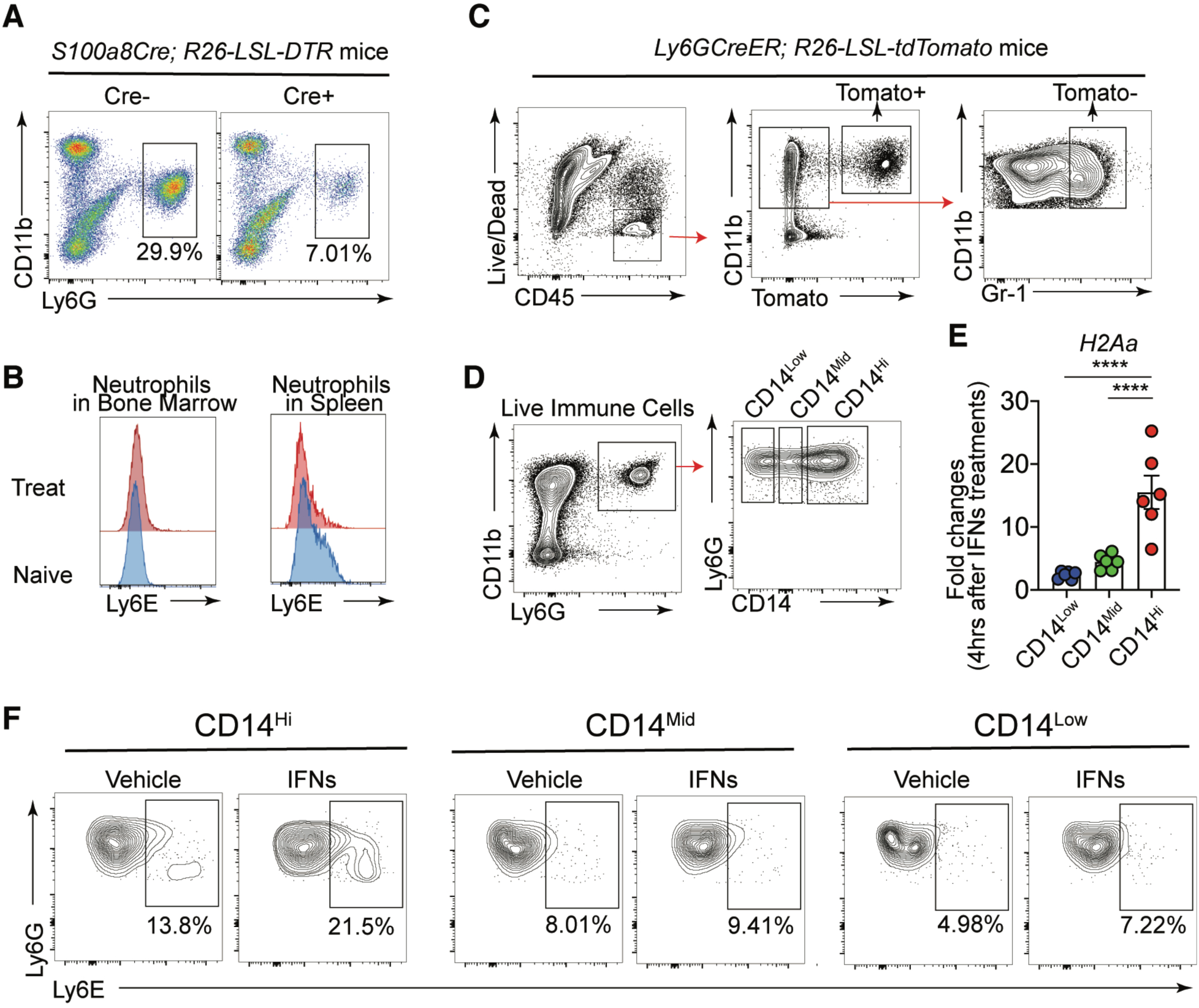
Flow cytometry analysis of dynamic changes in TANs during immunotherapy treatments. Related to Figure 2. **(A)** Representative flow cytometry plots showing efficient depletion of TANs after diphtheria toxin treatments of the tumors formed on the Neu^DTR^ mice. **(B)** Flow cytometry histogram showing the unchanged expression of markers associated with anti-tumor activity (e.g. Ly6E) in neutrophils in other tissues (bone marrow or spleen) from the same tumor-bearing mice after immunotherapy treatment. **(C)** Representative flow cytometry plots showing the sorting strategy to isolate lineage traced Tomato+ neutrophils that are present in the tumors before the immunotherapy treatments. **(D** to **F)** Representative flow cytometry plots (D) showing the sorting strategy to isolate CD14^Hi^, CD14^Mid^, and CD14^Low^ TANs for qPCR (E) and flow cytometry (F) to investigate how *in vitro* interferon treatments differentially reprogram various subsets of TANs.

**Fig. S3.**
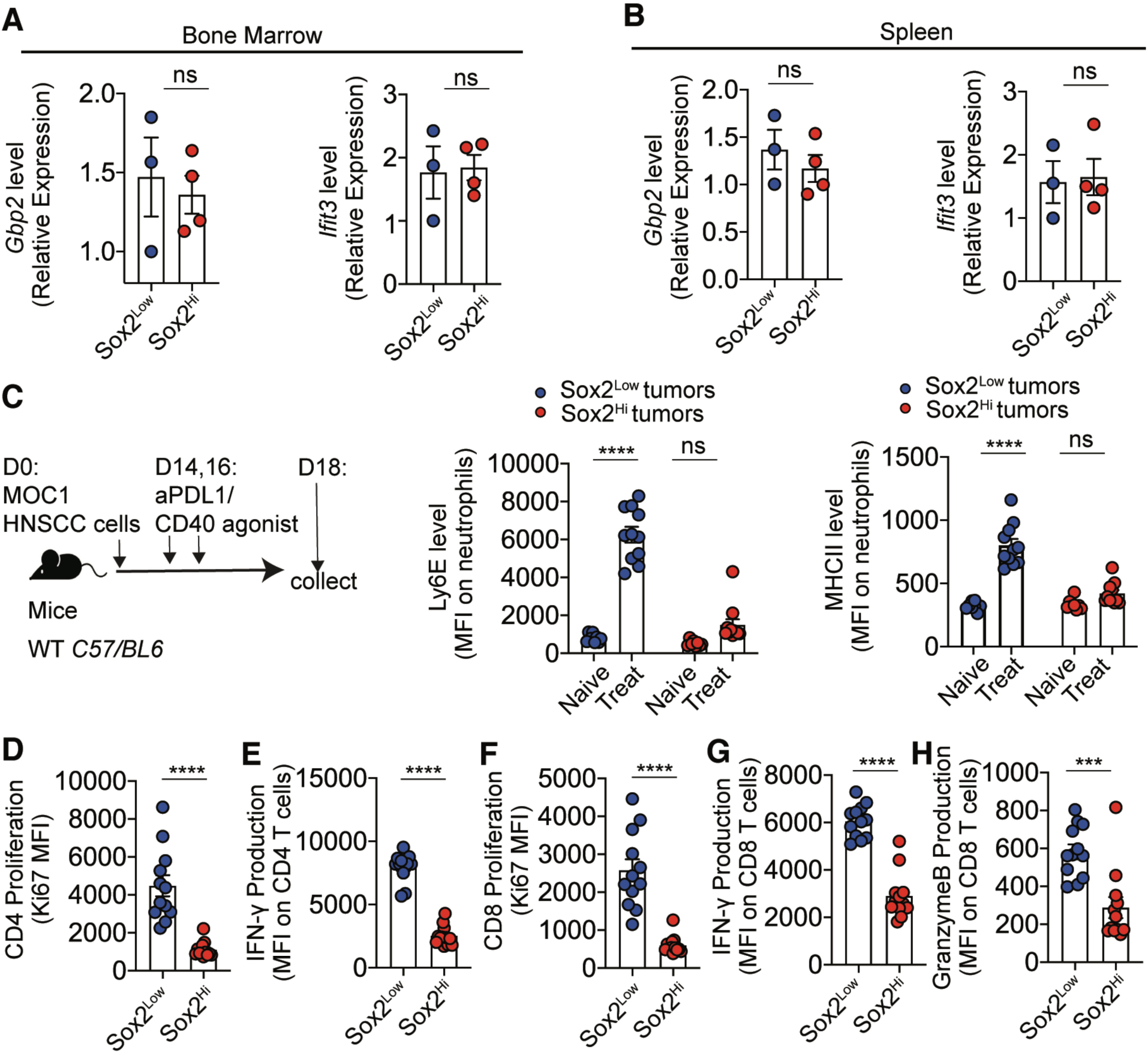
Immunotherapy induces distinct responses in different neutrophil subpopulations. **(A** and **B)** Quantitative PCR measuring the *Gbp2* and *Ifit3* expression in neutrophils isolated from bone marrow (A) or spleen (B) of the naive mice grafted with Sox2^Low^ or Sox2^Hi^ tumors following *in vitro* IFNα/IFNγ treatments for 4 hr. **(C)** Experimental scheme and flow cytometry quantification of the Ly6E and MHCII expression in TANs in HNSCC tumor formed by MOC1 cells with or without amplifying Sox2 before and after anti-PDL1/CD40 agonist treatments, n=10 in each group. **(D to H)** Flow cytometry quantification of CD4 T cell proliferation (D) and IFNγ production (E), as well as CD8 T cell proliferation (F), IFNγ (G) and Granzyme (H) production in HNSCC tumors formed by MOC1 cells with or without amplifying Sox2. n=12 in each group. Bar graphs show representative results from one the two repeats for each experiment and are presented as mean ± SEM. **** p < 0.0001; ns, non-significant.

**Table S1.**
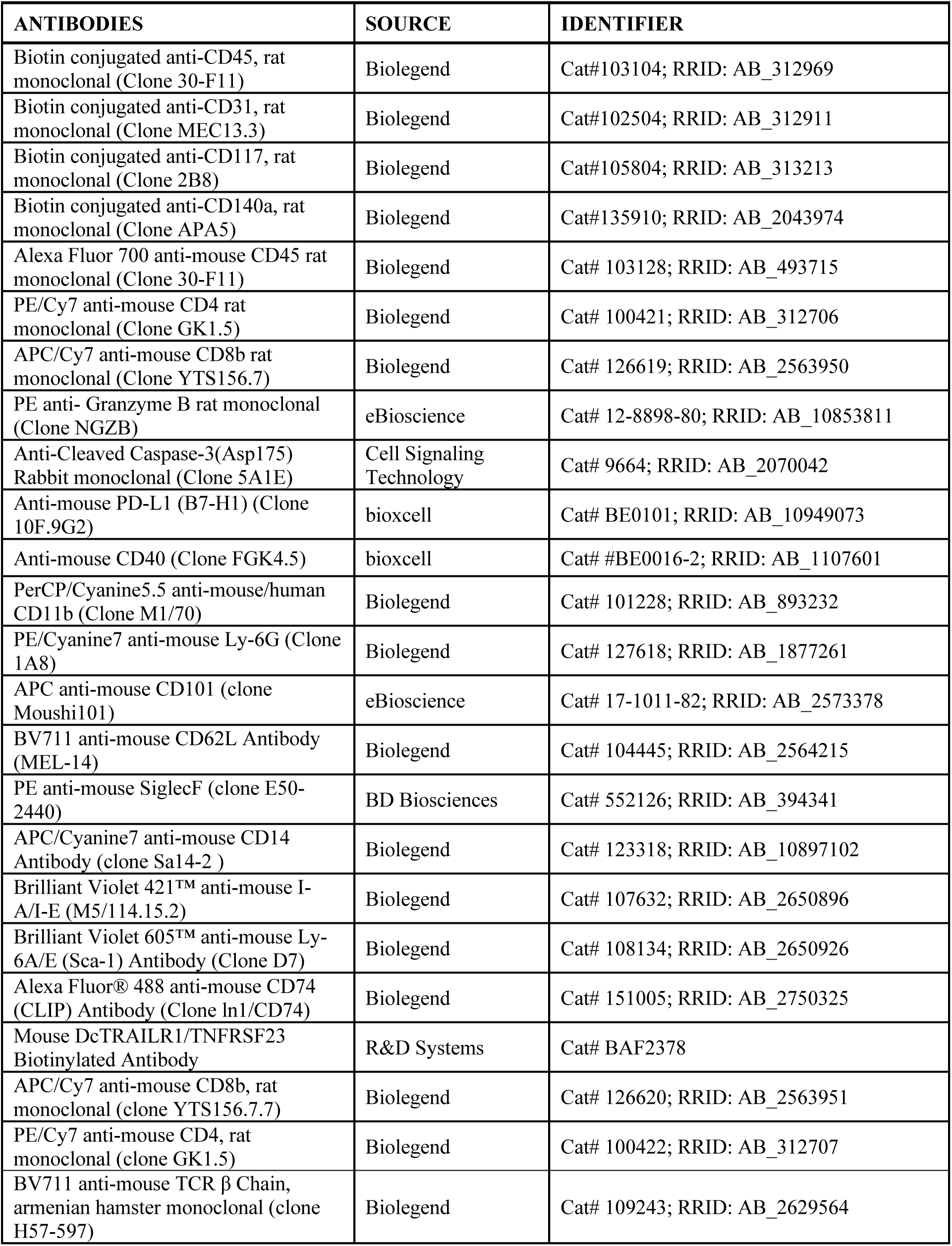
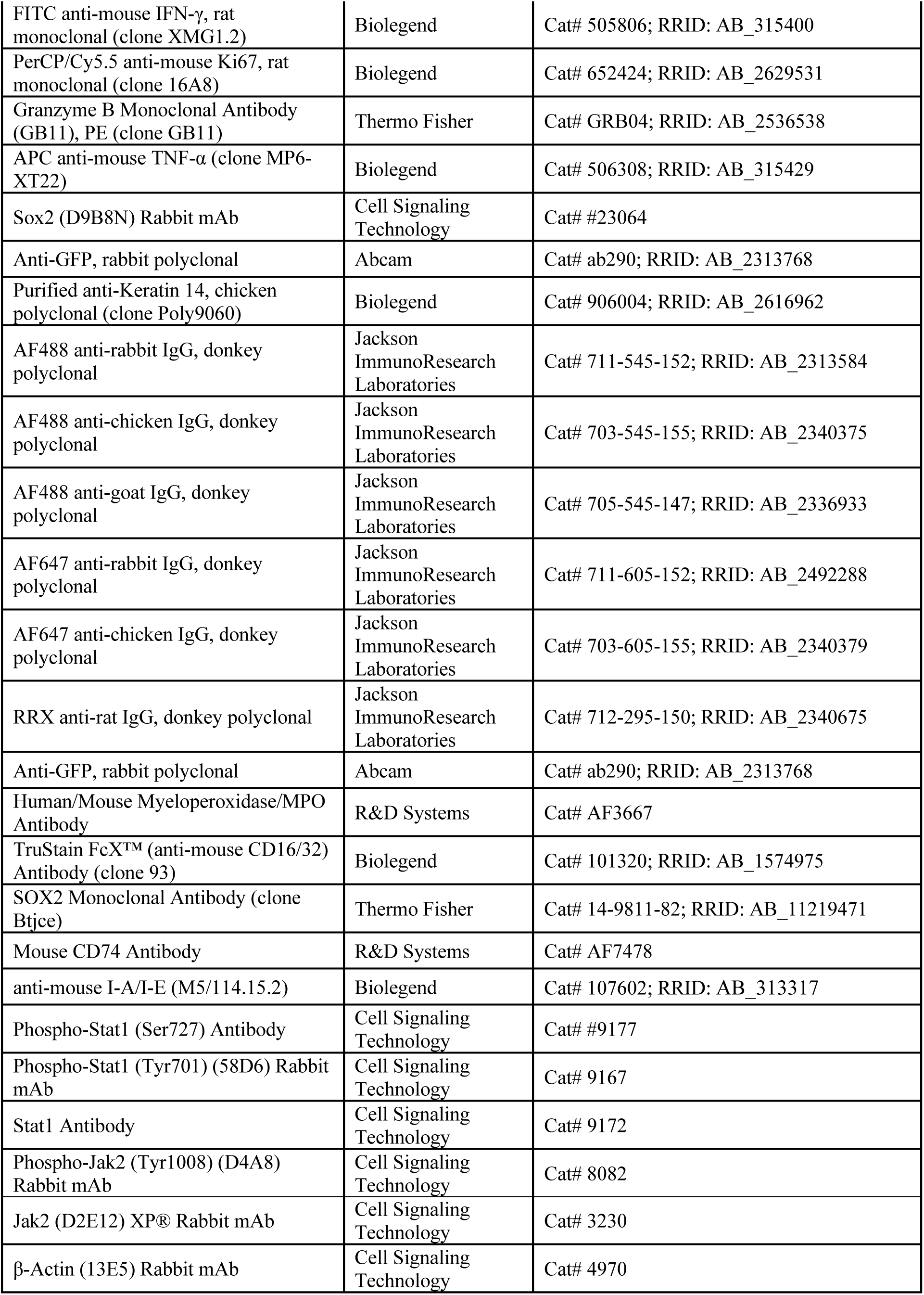
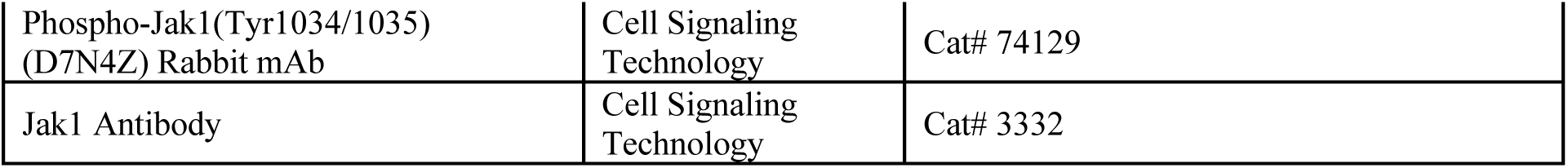
List of Antibodies.

**Table S2.**
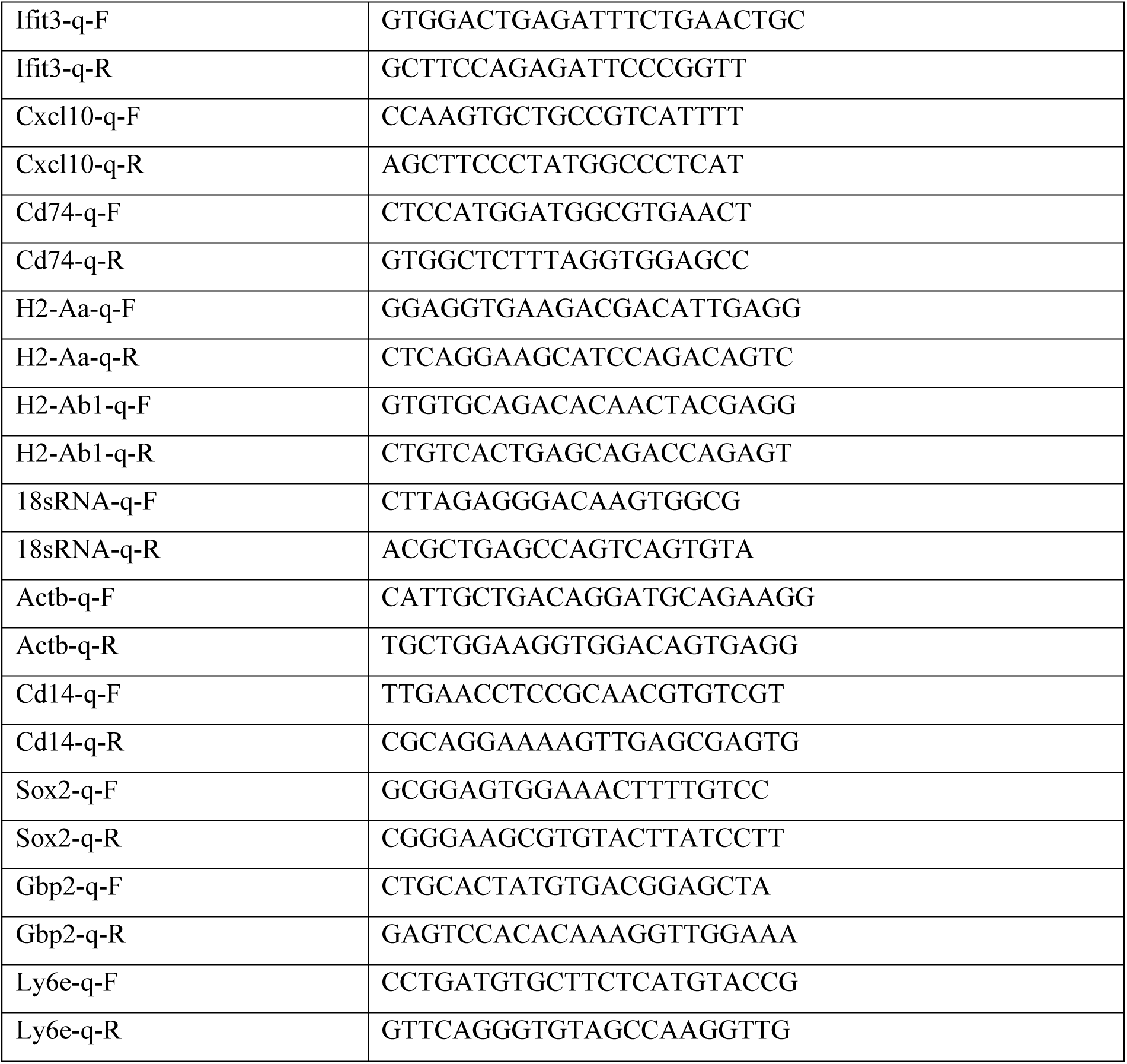
List of Quantitative PCR (qPCR) Primers.

**Table S3.** List of Reagents.

| REAGENT | SOURCE | IDENTIFIER |
| --- | --- | --- |
| Tamoxifen | Sigma | Cat# T5648 |
| Diphtheria toxin | Sigma | Cat# D0564 |
| Cultrex BME, Type 3 gels | Bio-Techne | Cat# 3632-005-02 |
| arachidonic acid | MP BIOMEDICALS | Cat# 1713889 |
| beta-mercaptoethanol | GIBCO | Cat# 21985023 |
| insulin | Sigma | Cat# I6634 |
| Hydrocortisone | Sigma | Cat# H0135 |
| Epidermal Growth Factor (EGF), human recombinant | Sigma | Cat# 01-107 |
| Recombinant Mouse GM-CSF | Biolegend | Cat# 576306 |
| Recombinant Mouse IFN-α | Biolegend | Cat# 752802 |
| Recombinant Mouse IFN-γ | Biolegend | Cat# 575302 |
| RNAscope Probe-Mm-Gbp2-C2 | Advanced Cell Diagnostics | Cat# 572491-C2 |

## Notes

### Competing Interest Statement

The authors have declared no competing interest.

## Reference

1. A. Ribas, J. D. Wolchok, Cancer immunotherapy using checkpoint blockade. Science 359, 1350–1355 (2018).

2. L. Pala et al., Outcomes of patients with advanced solid tumors who discontinued immune-checkpoint inhibitors: a systematic review and meta-analysis. EClinicalMedicine 73, 102681 (2024).

3. A. Betof Warner et al., Long-Term Outcomes and Responses to Retreatment in Patients With Melanoma Treated With PD-1 Blockade. J Clin Oncol 38, 1655–1663 (2020).

4. A. S. Patel, I. Yanai, A developmental constraint model of cancer cell states and tumor heterogeneity. Cell 187, 2907–2918 (2024).

5. D. Nassar, C. Blanpain, Cancer Stem Cells: Basic Concepts and Therapeutic Implications. Annu Rev Pathol 11, 47–76 (2016).

6. J. J. Loh, S. Ma, Hallmarks of cancer stemness. Cell Stem Cell 31, 617–639 (2024).

7. E. Batlle, H. Clevers, Cancer stem cells revisited. Nat Med 23, 1124–1134 (2017).

8. P. Baldominos et al., Quiescent cancer cells resist T cell attack by forming an immunosuppressive niche. Cell 185, 1694–1708 e1619 (2022).

9. Y. Miao et al., Adaptive Immune Resistance Emerges from Tumor-Initiating Stem Cells. Cell 177, 1172–1186 e1114 (2019).

10. A. Miranda et al., Cancer stemness, intratumoral heterogeneity, and immune response across cancers. Proc Natl Acad Sci U S A 116, 9020–9029 (2019).

11. R. E. Niec, A. Y. Rudensky, E. Fuchs, Inflammatory adaptation in barrier tissues. Cell 184, 3361–3375 (2021).

12. S. Naik, S. B. Larsen, C. J. Cowley, E. Fuchs, Two to Tango: Dialog between Immunity and Stem Cells in Health and Disease. Cell 175, 908–920 (2018).

13. C. Liu et al., Niche inflammatory signals control oscillating mammary regeneration and protect stem cells from cytotoxic stress. Cell Stem Cell 31, 89–105 e106 (2024).

14. J. Fujisaki et al., In vivo imaging of Treg cells providing immune privilege to the haematopoietic stem-cell niche. Nature 474, 216–219 (2011).

15. D. Bayik, J. D. Lathia, Cancer stem cell-immune cell crosstalk in tumour progression. Nat Rev Cancer 21, 526–536 (2021).

16. M. S. F. Ng et al., Deterministic reprogramming of neutrophils within tumors. Science 383, eadf6493 (2024).

17. C. Cui et al., Neutrophil elastase selectively kills cancer cells and attenuates tumorigenesis. Cell 184, 3163–3177 e3121 (2021).

18. D. Hirschhorn et al., T cell immunotherapies engage neutrophils to eliminate tumor antigen escape variants. Cell 186, 1432–1447 e1417 (2023).

19. P. Li et al., Dual roles of neutrophils in metastatic colonization are governed by the host NK cell status. Nat Commun 11, 4387 (2020).

20. Y. Wu et al., Neutrophil profiling illuminates anti-tumor antigen-presenting potency. Cell 187, 1422–1439 e1424 (2024).

21. S. Singhal et al., Origin and Role of a Subset of Tumor-Associated Neutrophils with Antigen-Presenting Cell Features in Early-Stage Human Lung Cancer. Cancer Cell 30, 120–135 (2016).

22. M. Benguigui et al., Interferon-stimulated neutrophils as a predictor of immunotherapy response. Cancer Cell 42, 253–265 e212 (2024).

23. J. Gungabeesoon et al., A neutrophil response linked to tumor control in immunotherapy. Cell 186, 1448–1464 e1420 (2023).

24. S. Jaillon et al., Neutrophil diversity and plasticity in tumour progression and therapy. Nat Rev Cancer 20, 485–503 (2020).

25. M. Quintanilla, K. Brown, M. Ramsden, A. Balmain, Carcinogen-specific mutation and amplification of Ha-ras during mouse skin carcinogenesis. Nature 322, 78–80 (1986).

26. D. Nassar, M. Latil, B. Boeckx, D. Lambrechts, C. Blanpain, Genomic landscape of carcinogen-induced and genetically induced mouse skin squamous cell carcinoma. Nat Med 21, 946–954 (2015).

27. I. L. Linde et al., Neutrophil-activating therapy for the treatment of cancer. Cancer Cell 41, 356–372 e310 (2023).

28. A. B. Rosenberg et al., Single-cell profiling of the developing mouse brain and spinal cord with split-pool barcoding. Science 360, 176–182 (2018).

29. F. Veglia et al., Analysis of classical neutrophils and polymorphonuclear myeloid-derived suppressor cells in cancer patients and tumor-bearing mice. J Exp Med 218, (2021).

30. F. Veglia et al., Fatty acid transport protein 2 reprograms neutrophils in cancer. Nature 569, 73–78 (2019).

31. R. E. Menjivar et al., Arginase 1 is a key driver of immune suppression in pancreatic cancer. Elife 12, (2023).

32. A. Sadik et al., IL4I1 Is a Metabolic Immune Checkpoint that Activates the AHR and Promotes Tumor Progression. Cell 182, 1252–1270 e1234 (2020).

33. M. Rapp et al., CCL22 controls immunity by promoting regulatory T cell communication with dendritic cells in lymph nodes. J Exp Med 216, 1170–1181 (2019).

34. J. Teishima et al., Fibroblast Growth Factor Family in the Progression of Prostate Cancer. J Clin Med 8, (2019).

35. I. Ballesteros et al., Co-option of Neutrophil Fates by Tissue Environments. Cell 183, 1282–1297 e1218 (2020).

36. A. Hasenberg et al., Catchup: a mouse model for imaging-based tracking and modulation of neutrophil granulocytes. Nat Methods 12, 445–452 (2015).

37. N. Oshimori, D. Oristian, E. Fuchs, TGF-beta promotes heterogeneity and drug resistance in squamous cell carcinoma. Cell 160, 963–976 (2015).

38. M. Schober, E. Fuchs, Tumor-initiating stem cells of squamous cell carcinomas and their control by TGF-beta and integrin/focal adhesion kinase (FAK) signaling. Proc Natl Acad Sci U S A 108, 10544–10549 (2011).

39. A. J. Bass et al., SOX2 is an amplified lineage-survival oncogene in lung and esophageal squamous cell carcinomas. Nat Genet 41, 1238–1242 (2009).

40. K. Arnold et al., Sox2(+) adult stem and progenitor cells are important for tissue regeneration and survival of mice. Cell Stem Cell 9, 317–329 (2011).

41. A. Sarkar, K. Hochedlinger, The sox family of transcription factors: versatile regulators of stem and progenitor cell fate. Cell Stem Cell 12, 15–30 (2013).

42. S. Boumahdi et al., SOX2 controls tumour initiation and cancer stem-cell functions in squamous-cell carcinoma. Nature 511, 246–250 (2014).

43. J. M. Siegle et al., SOX2 is a cancer-specific regulator of tumour initiating potential in cutaneous squamous cell carcinoma. Nat Commun 5, 4511 (2014).

44. B. Koletzko et al., FADS1 and FADS2 Polymorphisms Modulate Fatty Acid Metabolism and Dietary Impact on Health. Annu Rev Nutr 39, 21–44 (2019).

45. B. Wang et al., Metabolism pathways of arachidonic acids: mechanisms and potential therapeutic targets. Signal Transduct Target Ther 6, 94 (2021).

46. T. Lammermann et al., Neutrophil swarms require LTB4 and integrins at sites of cell death in vivo. Nature 498, 371–375 (2013).

47. J. J. Apiz Saab, A. Muir, Tumor interstitial fluid analysis enables the study of microenvironment-cell interactions in cancers. Curr Opin Biotechnol 83, 102970 (2023).

48. H. Tallima, R. El Ridi, Arachidonic acid: Physiological roles and potential health benefits - A review. J Adv Res 11, 33–41 (2018).

49. J. N. Cohen, et al., Regulatory T cells in skin mediate immune privilege of the hair follicle stem cell niche. Sci Immunol 9, eadh0152 (2024).

50. L. V. Nguyen, R. Vanner, P. Dirks, C. J. Eaves, Cancer stem cells: an evolving concept. Nat Rev Cancer 12, 133–143 (2012).

51. F. Baharom et al., Systemic vaccination induces CD8(+) T cells and remodels the tumor microenvironment. Cell 185, 4317–4332 e4315 (2022).

52. A. M. Luoma et al., Tissue-resident memory and circulating T cells are early responders to pre-surgical cancer immunotherapy. Cell 185, 2918–2935 e2929 (2022).

53. G. Micevic et al., IL-7R licenses a population of epigenetically poised memory CD8(+) T cells with superior antitumor efficacy that are critical for melanoma memory. Proc Natl Acad Sci U S A 120, e2304319120 (2023).

